# The Predictive Global Neuronal Workspace: A Formal Active Inference Model of Visual Consciousness

**DOI:** 10.1101/2020.02.11.944611

**Authors:** Christopher J. Whyte, Ryan Smith

## Abstract

The global neuronal workspace (GNW) model has inspired over two decades of hypothesis driven research on the neural basis consciousness. However, recent studies have reported findings that are at odds with empirical predictions of the model. Further, the macro-anatomical focus of current GNW research has limited the specificity of predictions afforded by the model. In this paper we present a neurocomputational model – based on Active Inference – that captures central architectural elements of the GNW and is able to address these limitations. The resulting ‘predictive global workspace’ casts neuronal dynamics as approximating Bayesian inference, allowing precise, testable predictions at both the behavioural and neural levels of description. We report simulations demonstrating the model’s ability to reproduce: 1) the electrophysiological and behaviour results observed in previous studies of inattentional blindness; and 2) the previously introduced four-way taxonomy predicted by the GNW, which describes the relationship between consciousness, attention, and sensory signal strength. We then illustrate how our model can reconcile/explain (apparently) conflicting findings, extend the GNW taxonomy to include the influence of prior expectations, and inspire novel paradigms to test associated behavioural and neural predictions.

## 1. Introduction

Global workspace theory (GWT) is one of the most widely supported neuroscientific theories of consciousness (Michel et al., 2018). GWT was first proposed by Baars (1988) as a cognitive architecture that identifies consciousness with the global availability of information. According to GWT, information becomes conscious when it is simultaneously made available to a wide range of localized (and individually sub-personal) processes – jointly comprising a ‘global workspace’. More recently, Dehaene and colleagues have advanced a global *neuronal* workspace (GNW) model, which identifies the global workspace with a large-scale network of excitatory pyramidal neurons with long-range axonal pathways connecting prefrontal and parietal cortices (Dehaene & Changeux, 2011; Dehaene et al., 2011; Mashour, Roelfsema, Changeux, & Dehaene, 2020). The key working hypothesis of the GNW is that when a stimulus becomes conscious there will be a late, non-linear, all-or-nothing “ignition” of prefrontal and parietal regions (Dehaene and Changeux, 2011; Dehaene, 2014; Mashour et al., 2020) corresponding to the large-scale influence of selected (otherwise unconscious) representations of perceptual features encoded locally within sensory cortices. In contrast, activity related to stimuli that is rendered unconscious (e.g., by masking or inattention) will fail to attain this global influence and related neuronal activity will only be observable locally within sensory cortices.

Behavioural and neurobiological predictions of the GNW can be broadly summarised in terms of a four-way taxonomy describing the relationship between consciousness, bottom-up sensory signal strength, and attention-based modulation (Dehaene et al., 2006). Specifically, in the absence of attention the activation caused by the presence of a weak stimulus should remain within early extrastriate areas, leading to weak priming effects (i.e. only slightly above chance) and unavailability for conscious report. When stimulus strength is weak but attention is present, the signal should reach deeper levels of extrastriate cortex – leading to noticeable priming effects (i.e. above chance) yet still unavailable for report. When stimulus strength is increased, but attention is absent, this should allow for deep processing, again facilitating noticeable (e.g., semantic) priming effects but activation should remain limited to sensory areas and be unavailable for report. Finally, when a strong signal has made its way to deep levels of processing and is amplified by top-down attention, prefrontal and parietal loops will be recruited to maintain sensory information through recurrent activity – thereby making it broadly available to large-scale networks subserving domain-general (goal-directed) cognition and allowing for conscious report (among other adaptive uses).

These predictions, while by no means uncontroversial, have been largely corroborated. In a pioneering fMRI study, Dehaene and colleagues found that conscious report of rapidly presented words resulted in the wide-spread activation of prefrontal, temporal and parietal regions, whereas activity remained within sensory regions when the stimulus was rendered invisible via masking (Dehaene et al, 2001). In a meta-contrast masking paradigm combined with electroencephologaphy (EEG), DelCul et al (2007) found that early event-related potential (ERP) components did not display a significant difference between seen and unseen conditions, while the late P3 component showed a significant non-linear increase in amplitude between seen and unseen conditions. Importantly, objective task performance was well above chance even at the lowest visibility rating. Similarly, using an attentional blink paradigm, Sergent et al (2005) found that early ERP components either did not vary with visibility, or had a linear relationship with visibility. In contrast, late components, such as the P3, showed non-linear increases in amplitude when T2 visibility was above 50%. Work using multivariate decoding in magnetoencephalography (MEG) has extended this finding, showing that, in comparison to temporally adjacent distractor stimuli, consciously reported target stimuli display a stronger and more sustained pattern of activation (Marti & Dehaene, 2017). Convincingly, when recording from intracranial electrodes implanted in epilepsy patients, Gaillard et al (2009, p.475) found that conscious word perception had a significant effect on many frontal and parietal electrodes, whilst the electrodes showing a significant effect of unconscious perception were almost all located in the occipital and temporal lobes.

Most recently, van Vugt et al, (2018) used multiunit electrodes implanted in non-human primate V1, V4, and dlPFC to study the relationship between the threshold for conscious access and ignition dynamics in PFC. Monkeys were trained to report a visual stimulus presented at low contrasts by making a saccade to a particular location. On unseen trials activity propagated through V1 (sometimes reaching V4), but the signal was lost before reaching PFC. In contrast, on seen trials activity was propagated with a high firing rate through V1 and V4, and caused a non-linear activation of dlPFC. Crucially, although not as pronounced as on seen trials, false alarm trials were also characterised by a spontaneous ignition like pattern of activation in dlPFC. These results support the conclusions of a recent modelling study (Joglekar et al, 2018), which showed that ignition like dynamics emerge naturally from a neural network constructed to mirror connectivity of the macaque cortex. As with van Vugt et al, (2018), they found that feedforward connections, when balanced by local inhibitory connections, steadily propagate activity forwards until the signal reaches PFC, at which point feedback connections cause a late and non-linear ignition like pattern of activity (for a discussion and up to date review of the theory and evidence behind the GNW see Mashour et al., 2020).

Despite the flood of research supporting the GNW, the theory also has a number of limitations. First, it is largely described at the level of gross neuroanatomy, leaving the details of cortical micro-architecture unspecified, which in turn limits limiting the granularity of predictions. Second, the GNW is agnostic about the implementation of expectation, rendering the theory unable to engage with a large body of evidence highlighting the role of expectation in visual consciousness (e.g. Chang et al., 2015; Denison et al., 2011; Valuch & Kulke, 2019; van Gaal et al., 2015).

Of greater concern is the fact that, as experimental paradigms have become more sophisticated, two predictions of the original theory have been falsified. For example, in a delay matching task, multivariate decoding showed that the brain represented both target presence and target orientation for an entire 800ms delay period across visibility levels. In addition, whilst target visibility correlated with decoding accuracy for target presence, unseen stimuli still exhibited a stable pattern of activation that generalised across time, suggesting that information did not have to be conscious to enter later phases of processing (King et al.,2016). This corroborates the findings of Salti and colleagues (Salti et al.,2015) who showed that target position could be decoded from superior frontal and superior parietal cortices in both seen and unseen conditions. Together these results demonstrate that, contrary to the original formulation of the GNW, information is processed and unconsciously maintained (at least briefly and insofar as multivariate decoding is a defensible proxy for information processing) by the same structures implicated in conscious processing.

Another seemingly inconsistent result was found by Pitts et al (2014). Using a novel inattentional blindness paradigm, they showed that the P3, which was initially thought to discriminate whether information had entered the global workspace (see Dehaene, 2014, p.180), was driven by task relevance and not conscious access. This result was recently replicated in a standard masking paradigm (Cohen, et al., 2020) showing unequivocally that the P3 is related to task relevance and is not a necessary signature of conscious access.

To the credit of GNW theorists, the model has been revised to accommodate these findings. First, a revised computational model of subjective report has been proposed, which, in addition to frontoparietal activity, requires that a stimulus representation can be separated from a noise distribution (King & Dehaene, 2014). Second, the claim that the P3 is a specific marker of conscious access (Dehaene et al., 2014) is no longer defended. However, the modified computational model of subjective report (King & Dehaene, 2014) is too idealised to make neurobiological predictions. In addition, the abandonment of the P3 as a signature of conscious access was not accompanied by any theoretical revisions and makes no additional behavioural or neurobiological predictions. This situates the GNW in a tenuous position, in which revision primarily explains away contrary results – a recognized characteristic of degenerative research programs (Lakatos, 1970).

Here we aim to make progress in overcoming these limitations by advancing an extension of the global neuronal workspace - the *predictive* global neuronal workspace (PGNW) - that unifies essential aspects of the GNW with the more recent (Bayesian) Active Inference approach to understanding brain function. Specifically, we present a hierarchical, partially-observable Markov decision process (POMDP) model of visual consciousness based upon Active Inference. Importantly, we leverage the neural process theory associated with Active Inference to make explicit links between neurobiology and the simulations afforded by the model.

Formalising ideas first introduced by Hohwy (2013), Whyte (2019), and Friston et al (2012), we will argue that conscious access or “ignition” is a fundamentally inferential process that depends upon a level of processing of sufficient temporal depth to contextualise and coordinate lower levels of processing. This longer timescale coordination is seen as necessary for the generation of subjective reports. Here, subjective reports stand in as one example of a broader set of temporally extended action plans (i.e., extended sequences of actions), the generation of which requires integration, maintenance, and manipulation of information over sufficient lengths of time – and where that information is sufficiently complex to guide the controlled generation of such goal-directed behaviours. For instance, combining conceptual contents associated with words such as “I”, “see”, “a”, “red” and “square” requires representing contents of much greater abstraction and temporal depth than is necessary for representing the perceptual property denoted by the word “red”.

To build on the previous conceptual contributions in this area mentioned above, we substantiate our arguments with a series of detailed computational simulations. These simulations are based upon the first principles account of perception and action selection offered by Active Inference. The simulations we show are also implemented using standard routines (that are available via open access; see software note), which will allow the reader to replicate our results and customize these simulations for their own purposes. The proof of principle offered by these simulations is particularly important when it comes to understanding the neuronal processes that implement the belief updating underlying the GNW formulation of conscious access. We mention this here to prepare the reader for a somewhat lengthy paper that must first cover some fundamentals that may appear a bit technical. However, as we will show, having an *in silico* subject at hand allows us to demonstrate how a host of otherwise disparate findings in current neuroscience research on visual consciousness can be explained by a first principles account of brain function.

To begin, we provide a brief primer on Active Inference and POMDPs, followed by a specification of the specific structure of the generative model we will employ, paying close attention to the importance of temporal depth. With the groundwork laid out, we show through simulations that the model can both; 1) unify seemingly contradictory previous results; and 2) reproduce the essential aspects of the four-way taxonomy predicted by the GNW, describing the relationship between conscious access, attention, and stimulus strength. Using the same generative model architecture, we then reproduce (and offer mechanistic explanations for) the electrophysiological and behavioural results of the inattentional blindness paradigm introduced by Pitts et al (2014).

Next, we turn to the role of expectation in visual consciousness and show how our model can extend the original four-way taxonomy of GNW theory to encompass paradigms that manipulate prior expectations on a trial-by-trial basis – highlighting the novel predictions that emerge from this extension. We also describe a novel paradigm that could be used to test distinct model predictions regarding dissociable effects of expectation, attention, and stimulus strength. We end by examining the relationship between the PGNW and alternative models, and briefly address potential concerns about how phenomenal consciousness could plausibly be situated within our model. However, this paper is chiefly concerned with what Block (2005) terms “access consciousness” which is defined as the availability of information for verbal report, voluntary action, and executive processing. For brevity, we will use “conscious access” and “consciousness” interchangeably throughout the paper unless otherwise indicated.

## 2. Theory

### 2.1 A Primer on Active Inference

Active Inference, a corollary of the free energy principle (FEP), is a first principles approach to modelling (approximately) Bayes optimal behaviour (Friston; 2010; Friston et al, 2016, 2017a). The FEP starts from the tautology that, in order for a self-organising system to maintain the integrity of its internal milieu, it must stay within the narrow range of states consistent with its survival. Human body temperature, for example, should ideally stay within the range of 36.5 – 37.5 degrees (Celsius). This entails that an organism’s phenotype has an attracting set of physical states. Over long timescales, these attractor states have a high probability of being observed in the sense that the organism will visit them repeatedly (Friston, 2013). Formally then, all self-organising systems must be minimising the (information-theoretic) surprise of their sensory observations (i.e., where observations deviating from phenotype-congruent states elicit surprise). However, surprise is computationally intractable.

Instead, according to the FEP, organism’s construct an internal (generative) model of environmental dynamics that, when accurate, acts as an upper bound on surprise (Buckley et al., 2017). The perception-action cycle is thus cast as an optimisation problem. Perception corresponds to the process of inferring the hidden state values that maximise the likelihood of observations and create a tight bound on surprise, while action is the process of inferring action sequences that either minimise uncertainty about hidden states (epistemic value) or bring about preferred observations (pragmatic value), thereby minimising the surprise following an action (Friston et al, 2016, 2017). The former (perception) corresponds to the minimisation of variational free energy **F**, while the latter (action selection) corresponds to the minimisation of expected free energy **G** (see figure 1 for formal descriptions).

**Figure 1.**
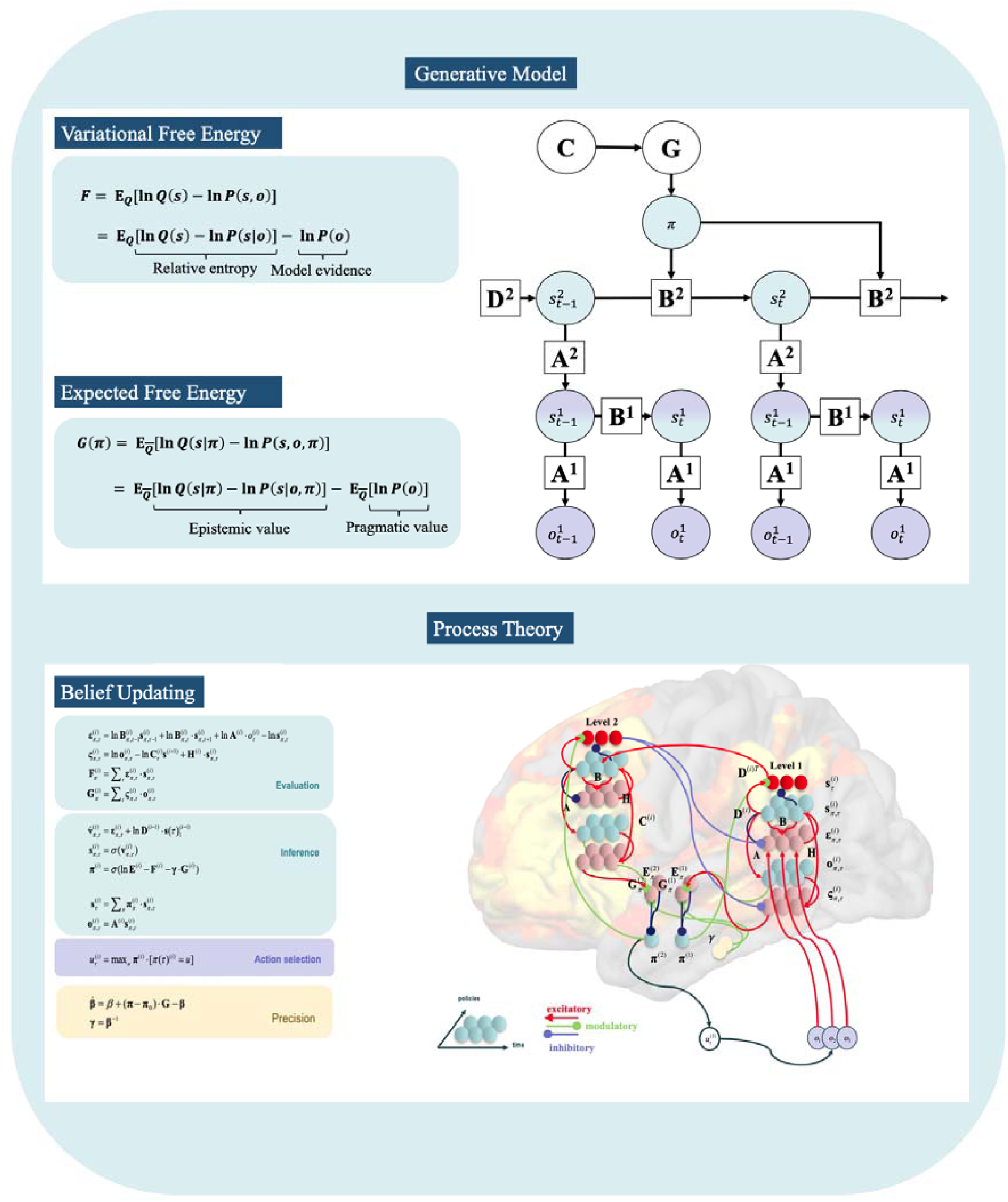
The **top panel** illustrates the free energy functionals and partially observable Markov decision process (POMDP) structure used within the generative model. The **left** side of this upper panel shows the decomposition of Variational free energy (VFE) into relative entropy and model evidence. Because the relative entropy term is always greater than or equal to zero when the approximate posterior approximates the true posterior VFE is equal to the negative model evidence. Minimising VFE is, therefore, equivalent to maximising model evidence. For visual simplicity we have not included the policy term in the VFE, although it should be noted that states are, generally, conditioned on policies. Also shown is the decomposition of expected free energy (**G**) into epistemic value and pragmatic value. Here 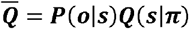 means that the expectation sums over all future observations that are expected under each policy (***π***). To minimise **G** agents must maximize the epistemic value term by selecting policies that transition them into states that maximising expected information gain (i.e. maximise the difference between **In *Q***(***s***| ***π***) and **In *P***(***s***|***o***, ***π***)) whilst also maximising the probability of preferred outcomes (**In *P***(***o***)) by seeking out preferred states. In other words, agents are driven to select actions that reduce uncertainty about hidden states whilst also aiming to maximise the fulfilment of their preferences. The **right** side of this upper panel provides a graphical depiction of the 2-level POMDP. Arrows show the dependences between variables (circles). Observations depend on hidden states at the first level. In turn, hidden states at the first level depend hidden states at the second level. Specifically, first-level hidden states function as observations for the second level. Equivalently, first level hidden states are the outcomes generated by hidden states at the second level. This representation also highlights the role of the vectors/matrices (squares) in determining the conditional dependencies between variables. Observations are generated by hidden states described by the matrix **A**. The **B** matrix determines state transitions, beliefs about which function as empirical priors. The **D** vector serves as the prior for initial states. When the **B** matrix is under the control of the agent, state transitions depend upon the policy (***π***). The probability that a particular policy will be selected depends on the expected free energy **G** of the policy which is, in turn, partially dependent on prior preferences specified by **C**. The **bottom panel** depicts the neural process theory associated with the model, Including the update equations and neural network implementation of the message passing implied by the Bayesian network shown in the top panel. The **left** portion of this lower panel shows the update equations and free energy functionals. Heuristically, state prediction errors ***ε*** score the evidence that observations provide for each policy (i.e., the difference between outcomes expected under each policy and those that are subsequently observed). In contrast, outcome prediction errors ***ς*** encode beliefs about the value of each policy (i.e., higher outcome prediction errors for a given policy roughly correspond to lower probabilities of observing preferred outcomes under that policy, as well as less informative observations expected under that policy). Directly below are expressions for variational free energy **F** and expected free energy **G**, expressed in terms of the above mentioned (state and outcome) prediction errors. Also shown are the update equations for states, policies, Bayesian model averages and the depolarisation variable, as well as action selection and its relation to the update terms for expected precision over policies. The **right** side of this lower panel provides a schematic of message passing between cell populations that could implement these updates. Red units encode Bayesian model averages, cyan units encode expectations over hidden states, and pink units encode state and outcome prediction errors. The backdrop image, depicting the neuroimaging signature associated with conscious access, is adapted from (Sadaghini et al., 2009).

Before we continue, it is worth acknowledging here that the material in this section might understandably come across to some readers as overly engineered and unnecessarily mathematical. However, at a computational level of analysis, the mechanics of belief updating described here are necessary to quantitatively account for both perception and action selection. Crucially, as we will see later, many aspects of the neuronal implementation of this belief updating scheme are already well established (at least at coarser-grained levels of description) and provide novel predictions in terms of firing rates and associated measures of synaptic efficacy.

Here we formulate the generative model as a partially observable Markov decision process (POMDP; see figure 1). POMDPs model discrete transitions between latent variables and the observations they generate. Such models infer states and policies based upon the (likelihood) mapping between different hidden state factors and distinct observation (or outcome) modalities – given by a set of **A** matrices (one matrix per outcome modality). Transitions between states are determined by the transition probabilities encoded by a set of **B** matrices (*at least* one matrix per state factor; see description of policy selection below). A set of **C** matrices describes the agent’s prior preferences over observations at each time point (one matrix for each outcome modality) and quantifies the degree to which agents prefer, or are averse to, particular observations. Finally, prior beliefs about initial states are determined by a set of **D** vectors (one per hidden state factor). **A, B, C** and **D** are each categorical distributions with Dirichlet priors over their respective parameter spaces.

Such models are equipped with allowable sequences of actions that can be chosen (plans or policies; π), where each possible sequence is assigned a value (higher policy values relate to lower expected free energies **G**, defined in relation to the prior preferences encoded in **C**). In the context of this class of models, allowable policies are specified as sequences of allowable state transitions, where each allowable transition (action) at each time point is encoded by a distinct **B** matrix for a given state factor. Thus, action corresponds to the agent’s direct control of state transitions. Observations and hidden states are factorised into separate outcome *modalities* and hidden state *factors* to allow for interactions between hidden states in the likelihood mapping (**A**). In hierarchical models, such as the model employed in this paper, the hidden states at the first level serve as observations at the second level (see figure 1). Crucially, hierarchical models also allow for inferences about deep temporal structure. An intuitive example of this is reading, in which the first level of a model could infer single words while the second level could infer the narrative meaning entailed by sequences of words over longer spatiotemporal scales (see Friston et al, 2017). Over the timescale of a single trial of a task (for example) belief updates are equivalent to (e.g., perceptual) *inference*, while, over longer timescales, updating gives rise to *learning* (we refer mathematically interested readers to Da Costa et al., 2020a). Technically, inference refers to updating beliefs about hidden states, while learning corresponds to updating (beliefs about) the parameters of the generative model specified by the matrices described above.

In terms of neurobiological implementation, Active Inference has a detailed process theory that specifies how a family of possible message passing algorithms can be used to perform inference, as implemented within neurobiologically plausible structural and functional dynamics (Friston et al., 2017; Parr & Friston, 2018).

Broadly speaking, the firing rates of certain neuronal populations - represented in figure 1 as neuronal populations within cortical columns - encode the current estimate of the posterior probability over hidden states. The synaptic inputs to the columns carry the conditional probabilities encoded in each of the matrices described above. This means that, for example, activity levels in neuronal populations encoding expectations of states are updated by ascending signals (from observations) based on synaptic weights encoding the amount of evidence that each possible observation provides for each possible hidden state (i.e., entries within the **A** matrices).

Of particular importance for the purposes of this paper are the equations describing the expected hidden states and the time derivative of the depolarisation variable 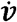 (see lower half of figure 1) associated with the neural process theory linked to Active Inference. Specifically, the posterior expectation over hidden states is a softmax (normalized exponential) function of the depolarisation variable *υ* which represents the average membrane potential of the neuronal populations responsible for encoding the surprise of expected states. The output of the softmax function is taken as the average firing rate of the population. The use of the softmax function (which is simply a generalisation of the sigmoid logistic function to vector inputs) to simulate average firing rate is based on the assumption made in mean-field models that the average firing rate of a population can be treated as a sigmoid function of the average membrane potential (Breakspear, 2017; Da Costa et al, 2020b). ERPs and local field potentials are modelled as the time derivative (rate of change) of the depolarisation variable because the change in membrane potential over neuronal populations is what generates ERPs. It is worth highlighting the face validity of this setup. Because the depolarisation variable is not normalised it can take both positive and negative values (i.e. like voltage) and after being normalised by the softmax function it is bounded between zero and one (i.e. like a normalised firing rate).

### 2.2 A Deep Temporal Model of Visual Consciousness

To model the difference between conscious and unconscious perception, we based our simulated task on the paradigm introduced by Pitts and colleagues (2012, 2014a, 2014b). We chose this task because, with only minor changes in design, the paradigm can be used to study both inattentional blindness and phenomenal masking – allowing us to model the interaction between attention and sensory signal strength in an empirically plausible manner.

At the beginning of each trial in our simulated task, the *in silico* subject - or agent - was presented with a stimulus composed of an array of bars surrounded by coloured disks. At the 2nd time point, the array of bars was replaced by a square, and at the 3rd time point the array changed back to the collection of bars. The agent was then required to self-report whether or not they had seen the square or to perform a two-alternative forced-choice task. We manipulated attention by requiring the agent to monitor the colour of the surrounding circles (either red or black) at the expense of the inner array (see figure 2).

**Figure 2.**
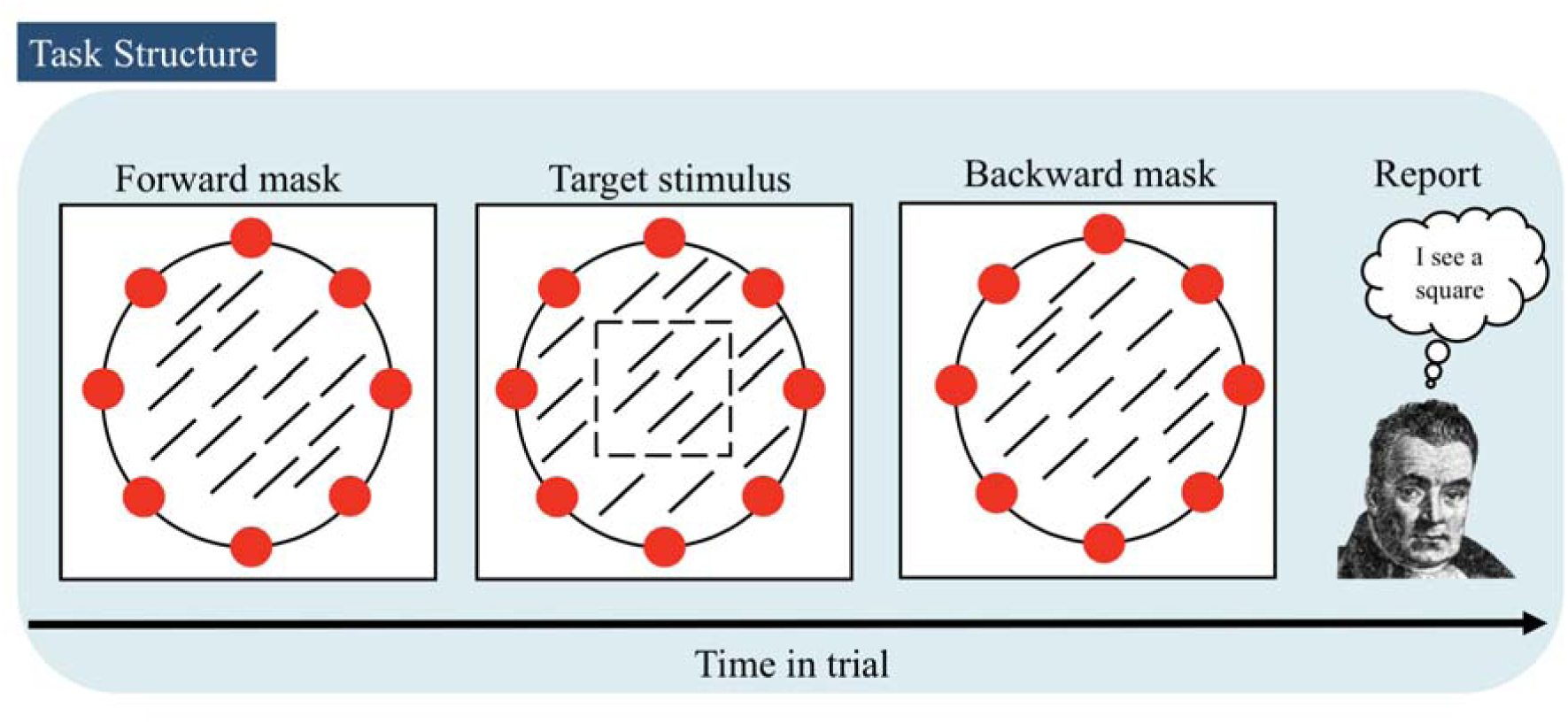
Illustration of the task performed by the agent. On each trial the in silico subject was presented with a stimulus composed of an array of bars surrounded by coloured discs. At the 2nd time point, the array was replaced by a square, and at the3rd time point the array changed back to the original bar pattern. The agent was then required to either perform a two-alternative forced-choice task or report whether they had seen the square.

To model manipulations of attention and stimulus signal strength, we manipulated the precision of the mapping between hidden states and the outcomes encoded in the first-level **A** matrix by passing what were initially identity matrices through two softmax (normalized exponential) functions controlled by two precision parameters, and, respectively encoding attention and signal strength (i.e. presentation time or contrast; see figure A1 in the appendix). Higher values of these parameters made the **A** matrices more precise. We set up the interactions between the **A** matrices such that the likelihood mapping for the internal stimulus factor was more precise when the agent was in an attentive state. In contrast, stimulus strength manipulations reduced the precision of the mapping between stimuli and hidden states independent of attentional state, with high stimulus strength corresponding to a precise likelihood mapping.

To simulate the perceptual categorisation and self-report behaviour described above, we specified the task in terms of a generative model which was inverted using a variational message passing scheme (for technical details see Parr et al., 2019a). To reproduce the recurrent interactions between the frontoparietal network identified with the global workspace, and the input it receives from the visual system, we employed a two-level deep temporal model (Friston et al., 2017b; Friston et al., 2018). The first level (see figure 3), which roughly corresponds to processes within the visual system, had four hidden state factors; attention allocation, internal stimulus (bars/square), surrounding or peripheral stimulus (red/black circles), and a set of auditory-verbal states (a number of words that could be put together in different sequences to generate verbal reports). The second, higher level, which corresponds to the frontoparietal network associated with the global workspace, included three hidden state factors: 1) sequence type (encoding beliefs about the sequence of internal and peripheral stimuli presented on each trial), 2) time point within the trial which, in line with data from non-human primates (Kapoor et al., 2018), encodes the current phase of the task, and 3) abstract semantic representations that could be unpacked into different verbally reported sequences of words at the lower-level (dependent on the chosen policy at the higher level).

**Figure 3.**
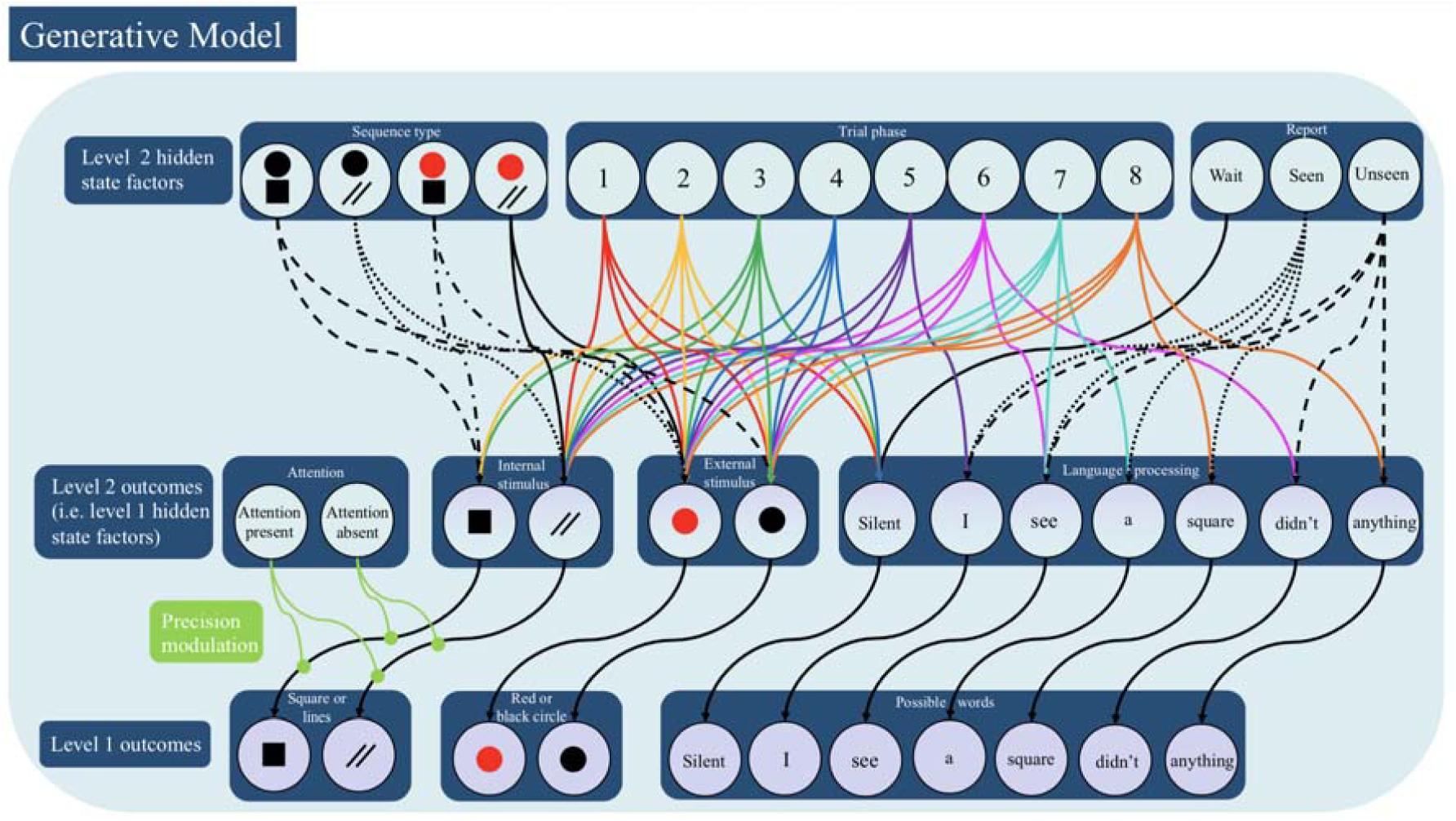
Bayesian network depiction of the generative model, with arrows showing the dependencies between hidden state factors and outcome modalities. At the second level, states within the sequence type and trial phase hidden state factors determine the internal stimulus and peripheral stimulus hidden states at the first level (which function as second-level observations). This was set up such that the mapping between second-level hidden states and first-level hidden states was dependent on the time point within the trial. The report state factor determined the language processing state at the first level. In turn, first-level outcomes were dependent on first-level hidden states. Importantly, there was only a precise mapping between the first-level internal (line/square) stimulus hidden state factor and the corresponding outcome modality when the model was in an attentive state.

It is worth emphasising that the temporal depth of this second level is essential for simulating the self-report behaviour that defines conscious access. The language component of the model is obviously an oversimplified depiction of linguistic cognition. Yet, it remains true that, in order to coordinate the selection of a specific sequence of words (i.e., to construct a sentence describing the content of perception), where each word is generated over a more rapid time scale, the agent *must* have a level of processing that unfolds over a slow enough time scale. This allows the agent to abstract away from the moment-to-moment sensory flux and coordinate lower-level language processes to report the outcome of perceptual decision-making in a goal-(i.e., preference-) dependant manner. Thus, the higher level is necessary to accumulate evidence from the moment-to-moment sensory flux with the longer-timescale, controlled processes capable of generating goal-directed behaviour plans with greater temporal depth. This feature is at the core of our model and we will revisit it in more depth in the discussion.

We made the modelling decision to have a more liberal threshold for forced-choice behaviour than for subjective reports based on the well-replicated finding that subject’s display above-chance performance even in the absence of reportability (see discussion in King & Dehaene, 2014). However, we acknowledge that this finding is largely dependent on the method of report. Specifically, there is evidence suggesting that humans have optimal introspective access to their perceptual processes in the sense that betting performance in a 2-interval forced-choice task matches that of an ideal Bayesian observer (Peters & Lau, 2015). We do not wish to take a stand on this issue here as we consider it an open empirical question.

Instead, we merely note that the decision was a pragmatic one based on the method of report used in the paradigms we were aiming to simulate and that we plan to pursue a more principled justification for report thresholds in future work.

For a full description of the generative model in terms of the matrices and associated parameter values see the appendix following the discussion section.

## 3. Results

### 3.1 Foundational Simulations

As a proof of principle, we simulated 200 trials, 100 of which were “square-present” trials. The model was in an attentive state (Σ = 1) and the stimulus also had a high stimulus strength (ζ = 1), corresponding to a long presentation time that enabled evidence accumulation resulting in a relatively precise hidden state-to-outcome mapping. We found that the model performed with 100% accuracy on the forced-choice task and when reporting the presence – and absence – of the square (see figure 4).

**Figure 4.**
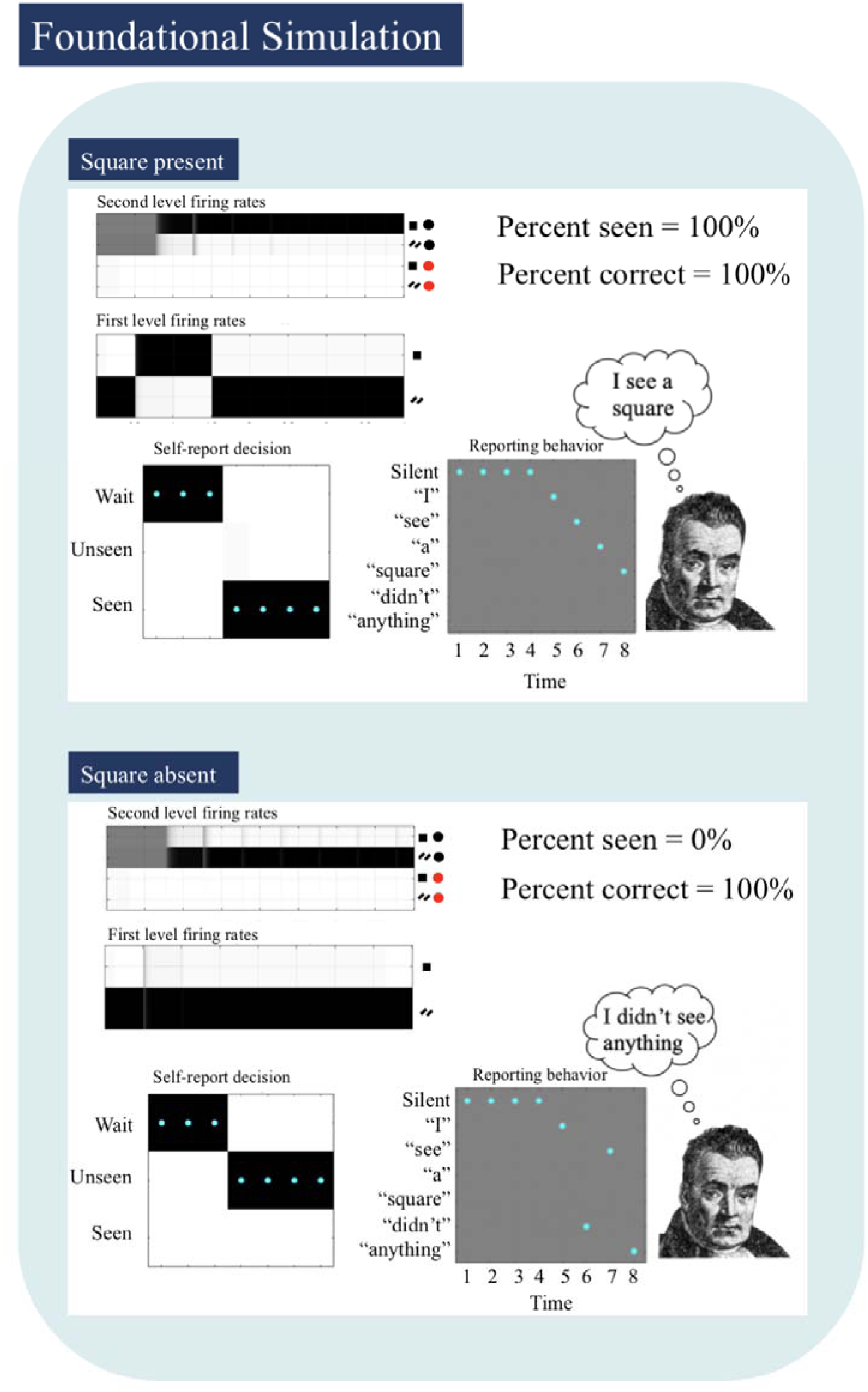
Simulated firing rates (darker = higher firing rate) predicted under the process theory associated with Active Inference (Friston et al., 2017). Each row represents the firing rate of the neuronal populations encoding the posterior expectations over each state. Individual squares each represent the time point within the trial. Actions (cyan dots = true action chosen; colour represents the posterior confidence in the actions chosen at the end of each trial, darker = higher confidence). Self-report outcomes (cyan dots = actual observations; colour indicates the preference for each observation, darker = greater preference, here the colours are all the same which means the agent did not have a preference for language outcomes). **Top**. Simulated results for 100 “square-present” trials. Notice the outcome modality associated with the “seen” report state generates the sentence “I see a square”. **Bottom**. Simulated results for 100 “square-absent” trials. Notice that the outcomes associated with the “unseen” report state generate the sentence “I didn’t see anything”.

Having established the face validity of the model, we now turn to the simulation of minimal contrast paradigms.

### 3.2 A Four-Way Taxonomy of the Factors Underlying Conscious Access

As a field, the neuroscience of consciousness has converged on the use of minimal contrast paradigms that, through masking, inattention, or near-threshold presentation, render nearly identical stimuli unconscious in one condition and conscious in the other. Combined with neuroimaging, this allows for the contrast of conscious and non-conscious forms of visual processing. However, early neuroimaging research reported conflicting results. Some studies found that subjective reports correlated with activation in early visual cortices (Zeki, 2003), while others found that subjective reports correlated with the activation of frontoparietal areas (for a meta-analysis see Bisenius et al., 2015). Still others argued that frontoparietal involvement is due to an attentional confound and does not reflect conscious access (Peter et al., 2005). Based upon simulations of thalamo-cortical networks (Dehaene et al., 2003), Dehaene and colleagues (Dehaene et al., 2006) created a taxonomy of factors underlying conscious access with the aim of unifying the conflicting results under the theoretical framework of the GNW.

In their exposition of the taxonomy, Dehaene and colleagues (Dehaene et al., 2006) distinguished between subliminal, preconscious and conscious forms of processing. However, for the sake of clarity, we describe the taxonomy in terms of the two factors underlying the classifications; attention, which can be present or absent, and signal (i.e. stimulus) strength, which can be strong or weak. As described in the introduction, the activation caused by a weak stimulus in the absence of attention (e.g. an unattended masked stimulus) should remain within early extrastriate regions – remaining unavailable for report and only causing weak priming effects. When attention is present, but stimulus strength is weak (an attended but masked stimulus), the signal should reach deeper levels of extrastriate cortex – remaining unavailable for report but leading to stronger priming effects.

When attention is absent, but stimulus strength is high (i.e. during inattentional blindness or motion-induced blindness), there should be deep levels of processing within sensory areas – facilitating priming effects but remaining unavailable for report. Lastly, when a strong signal reaches a deep level of processing and is amplified by top-down attention, recurrent loops in frontoparietal cortices will maintain the information over longer timescale and make it available to inform the generation of verbal reports and other long-timescale goal-directed behaviours.

Crucially, different neural correlates are observed when using specific paradigms, such as the attentional blink and phenomenal masking, to contrast different parts of the taxonomy. Dehaene et al (2006) have leveraged these findings to explain a number of seemingly contradictory results.

While the theoretical backbone of the taxonomy has been revised, in that the activation of frontoparietal regions is no longer considered sufficient for conscious access, it is still a useful starting point – as the interactions between stimulus strength and attention described by the taxonomy encompass many, if not most, of the minimal contrast paradigms reported in the literature.

As described above, we modelled the effects of both attention and signal strength on simulated behaviour by altering the precision of the state-outcome mapping between the internal stimulus hidden state and internal stimulus outcome at the first level. In terms of the task, the “low signal strength (ζ = 0.05) + attention absent (Σ = 0.01)” condition corresponds to a short presentation time, with the agent attending to the peripheral coloured disk at the expense of the internal stimulus. The “low signal strength (ζ = 0.01) + attention present (Σ= 1)” condition corresponds to a short presentation time with the agent directing attention to the internal stimulus.

The “high signal strength (ζ = 0.7) + attention absent (Σ = 0.1)” condition corresponds to a long presentation time with the agent attending to the peripheral coloured disk. Finally, the “high signal strength (ζ = 0.7) + attention present (Σ= 1)” condition corresponds to a long presentation time with the agent attending to the internal stimulus. We settled on the specific parameters shown above by searching the parameter space to find consistent values that best reproduced the behavioural results reported in the empirical literature, while remaining within plausible limits (i.e., as when fitting model parameters to real participant behaviour in empirical studies).

#### 3.2.1 Four-Way Taxonomy: Simulated Behaviour

We presented the model with 400 “square present” trials, 100 corresponding to each of the four parameter variants described above. When signal strength was low and attention absent, the agent displayed chance levels of forced-choice performance (47%) and only reported having seen the square on 10% of trials. When signal strength increased, or when the agent attended to the stimulus, forced choice performance improved to well above chance, 58% and 61% respectively, while still not reporting ‘seen’ on more than 40% of trials (11% and 36% respectively). When stimulus strength was strong, and attention was present, the agent showed near-ceiling levels of forced-choice performance (96%; i.e., due to the small amounts of stochasticity in choice within the model) and accurately reported having seen the square on 99% of the trials. Thus, under a consistent set of parameter settings, the model accurately reproduced the self-report and forced-choice behaviour commonly reported in the literature (see figure 5). This should not be a surprise, as we fine-tuned the parameter values to capture behaviour reported in the empirical literature. However, it is worth noting that it is possible a priori that no consistent set of parameter values could be found to reproduce all of these known empirical results – which would have shown a clear insufficiency of the model. Thus, the existence of this consistent set of parameter values does support the validity of the model.

**Figure 5.**
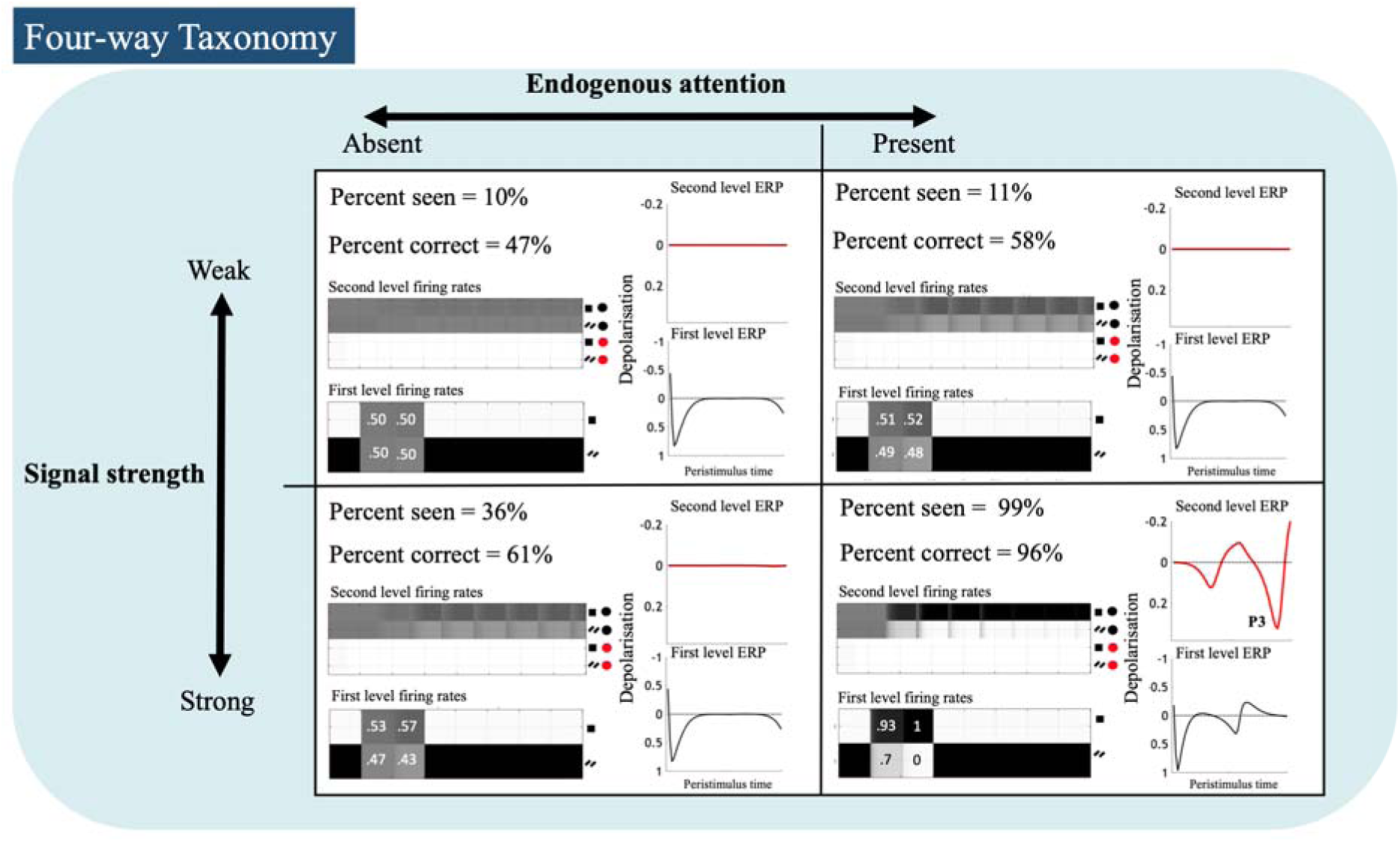
Report frequencies, forced choice accuracy, simulated firing rates, and simulated ERPs for each quadrant of the four-way taxonomy. Numbers shown on the lower-level firing rate plots illustrate the firing rate strengths (between 0 and 1) before and after top-down feedback from the higher level (i.e., time steps 2 and 3, respectively). ERP plots show the temporal derivative of the firing rates.

#### 3.2.2 Four-Way Taxonomy: Simulated Firing Rates

We next simulated the neural firing rates associated with the different quadrants of the taxonomy. The simulated first- and second-level firing rates both accurately reproduced a number of otherwise disparate empirical findings (see figure 5, left portion of each quadrant). Specifically, in all but the “weak signal + attention absent” condition, the model showed an amplification of first-level firing rates after receiving feedback from the second level. In line with the notion of ignition, this top-down amplification was notably greater when the model reported having seen the square. Similarly, at the second level of the model, in all but “weak signal + attention absent” condition the firing rates corresponding to the neuronal population encoding the “square present” sequence was amplified by the presentation of the square, even in the absence of report. Again, in line with the notion of ignition, the firing rate at the second level was further enhanced when the model reported having seen the square on all trials.

These results mirror the multivariate decoding results reviewed in the introduction, which found that, although increased visibility correlated with increased decoding accuracy, target orientation could be decoded across visibility levels from superior frontal and superior parietal cortices (Salti et al., 2015). What leads the agent to report the presence of the square is not simply that the second level has greater firing rates for the “square present” sequence, but that the model is sufficiently more confident about the “square present” sequence than the “random bar” (i.e. “square absent”) sequence. This feature of the model is similar to other Bayesian and hierarchical models in the literature that associate subjective report with the inference that a stimulus distribution is distinct from a noise distribution (King & Dehaene, 2014; Peters and Lau, 2015; Fleming 2019). However, the threshold for report in the present model is set by the expected free energy functional, which is composed of both an epistemic component (i.e., related to the difference between posterior expectations over states with and without an observation expected under each policy) and a pragmatic component (i.e., related to the deviation of expected and preferred observations under each policy; see Friston et al., 2017). The inclusion of the pragmatic component allows the PGNW to, in principle, explain the observed role of reward in modulating the content of visual consciousness (e.g. Marx & Einhäuser, 2015; Dong et al., 2019).

Importantly, if we were to apply a contrastive methodology and compare any of the three unconscious sections of the taxonomy with the conscious section, we would find that conscious access is associated with an enhanced firing rate at the first and second levels of the model. This mirrors the finding that conscious report correlates with the enhanced activation of sensory regions and the seemingly all-or-nothing activation of frontoparietal cortices originally cited in support of the GNW taxonomy (e.g. Dehaene et al, 2001). Based on our model, the (seemingly) contradictory finding that superior frontal and superior parietal cortices contain decodable information across visibility levels (Salti et al., 2015) can be explained as the result of using different methods to analyse the same underlying Active Inference architecture – that is, the use of standard univariate analyses on the one hand, which look for voxels that are more activated by visible stimuli than invisible stimuli (e.g. Dehaene et al., 2001), and more sensitive multivariate pattern analyses on the other, which instead look for patterns of information present across voxels. As was noted above, using a standard univariate approach to analyse the simulated firing rates shown in figure 5 would show that firing rate is enhanced on conscious compared to unconscious trials at both the first and second level. However, this ignores the presence of stimulus relevant information in the firing rates; although firing rates are higher for conscious compared to unconscious conditions, there is still a greater than baseline firing rate in unconscious conditions at both the first- and second-level within our simulations that could easily be exploited by a classifier. Thus, our model can simulate and explain both (seemingly conflicting) results within a single neurocomputational architecture.

#### 3.2.3 Four-Way Taxonomy: Simulated Event-Related Potentials

Finally, we examined the ERPs^1^ predicted by our model under different quadrants of the taxonomy (figure 5, right portion of each quadrant). Here we found that ERPs at the first level of the model – which correspond to early components such as the P100 and N100 – are relatively unaffected by changes in visibility. In contrast, ERPs at the second level – which correspond to late components, and specifically the P3 – appear to be strongly modulated by changes in visibility. This emulates the empirical findings described in the introduction, in which early components were relatively unaffected by changes in visibility while late components displayed a non-linear increase in amplitude as visibility increased (DelCul et al.,2007; Sergent et al., 2005).

As described above, ERPs are here modelled as the time derivative of the depolarisation variable. When confidence in a state changes rapidly, the derivative of the depolarisation variable increases in magnitude. This gives us a new perspective on late ERP components in the context of visual consciousness. Since the model only reports the presence of the square when it is sufficiently more confident about the “square present” sequence than the “random bar” sequence, it makes sense that verbal reports are accompanied by late ERPs – as they reflect the rapid change in beliefs at the second level of the model when the square is presented.

However, it is important to highlight that this is not a necessary feature of conscious access. If the model is already sufficiently confident in a state – either because of a precise likelihood mapping or because *the rate of belief updating is slowed* (by a lack of precision in the likelihood; e.g., because of inattention) – the model may report the presence of a stimulus without an accompanying late ERP. This is indeed what we see empirically. Specifically, in the inattentional blindness paradigm that guided development of the particular task structure of our model, Pitts and colleagues (2012, 2014a, 2014b) found that the P3 is associated with task relevance (i.e. attentional set) and *not reportability*, which was interpreted as evidence against the standard model of the GNW. In other words, the amplitude of the P3 changed as a function of attentional set – which affects precision – rather than conscious access as such. In the next section we discuss in more detail how our model can account for this result and demonstrate how the model can shed light on the specific electrophysiological correlates of inattentional blindness more generally.

### 3.3 Inattentional Blindness

The simulations reported in the previous section were focused on reproducing the findings that characterise minimal contrast paradigms associated with the GNW theory’s proposed four-way taxonomy. However, minimal contrast paradigms are not without limitations. For example, as Aru and colleagues (2012) have argued, these paradigms often confound the neural correlates of consciousness with the prerequisites to, and consequences of, consciousness. With these confounds in mind, in the study referred to above Pitts and colleagues (2014a, 2014b) created a three-phase sustained inattentional blindness paradigm that was designed to dissociate the electrophysiological correlates of consciousness from the correlates of task relevance.

The task used the same stimulus set discussed in the previous two sections.

In phase one, participants were instructed to monitor the peripheral disks for a change in colour. Every 600-800ms the internal section of the stimulus alternated between random bars and a square, both of which were presented for 300ms. After phase one, participants completed a debrief and 50% of them reported not being aware of the square, replicating the findings of Mack and Rock (1998). The debrief acted as a cue alerting the participants to the presence of the stimulus. Phase two was identical to phase one except that participants had been alerted to the presence of the square. Despite not being task relevant, 100% of participants reported having seen a square in the subsequent debrief. In phase three, participants were instructed to attend to the inside stimulus while ignoring the peripheral disks.

Contra the original predictions of the GNW model, the ERP results revealed a dissociation between awareness and the P3. As expected, in phase one the P3 was absent for inattentionally blind participants. However, the P3 was also absent in phase two when all participants were conscious of the stimulus despite it not being task relevant. In contrast, when the stimulus was conscious and task relevant (in phase three), there was a large P3, showing that task relevance is the primary diver of this ERP component.

Our aim in simulating this task here is to provide a bridge between established experimental findings and the more abstract account of the relationship between the P3 and visual consciousness advanced in the previous section. To match the empirical setup, we modelled phase one by placing the agent in an “attention absent” state toward the square stimulus (Σ = 0.05) and then found the value of the stimulus strength parameter (ζ= .75) that led the model to report the presence of the square ∼50% of the time (as during individual thresholding procedures in empirical studies). To model the effect of the debrief that alerted the participants to the presence of the square, in phase two we increased the value of the attention parameter (Σ = 0.2) so that it was higher than in the “attention absent” state but still substantially lower than an “attention present” state, plausibly corresponding to a diffuse allocation of attention. Phase three was the same as phase two, except that the model was placed in an “attention present” state toward the possibility of seeing the square (Σ = 1).

#### 3.3.1 Inattentional Blindness: Simulated Behaviour and Event Related Potentials

We presented the model with 300 square trials, 100 corresponding to each of the three sets of parameter values described above. Through stimulus strength calibration, in phase one the model reported the presence of the square 53% of the time, mirroring the empirical results. Crucially, however, the electrophysiological results of all three phases, and the behavioural results of phase two and phase three, reproduced the empirical findings without any further adjustments to parameter values. In phase one, there were only early first-level ERPs (see figure 6). In phase two, when the “square present” and “random bar” sequences had equal prior probability, the model reported the presence of the square on 99% of trials and, in line with empirical results, there were again only large first-level ERPs; no late P3-like ERPs were generated. Finally, in phase three, when the model was attending to the internal stimulus, the square was reported on 100% of trials and there was a large and late ERP at the second level resembling the P3. Considering the idealised nature of the model, the simulated ERPs displayed in figure 6 bear a striking resemblance to the empirically observed ERPs.

**Figure 6.**
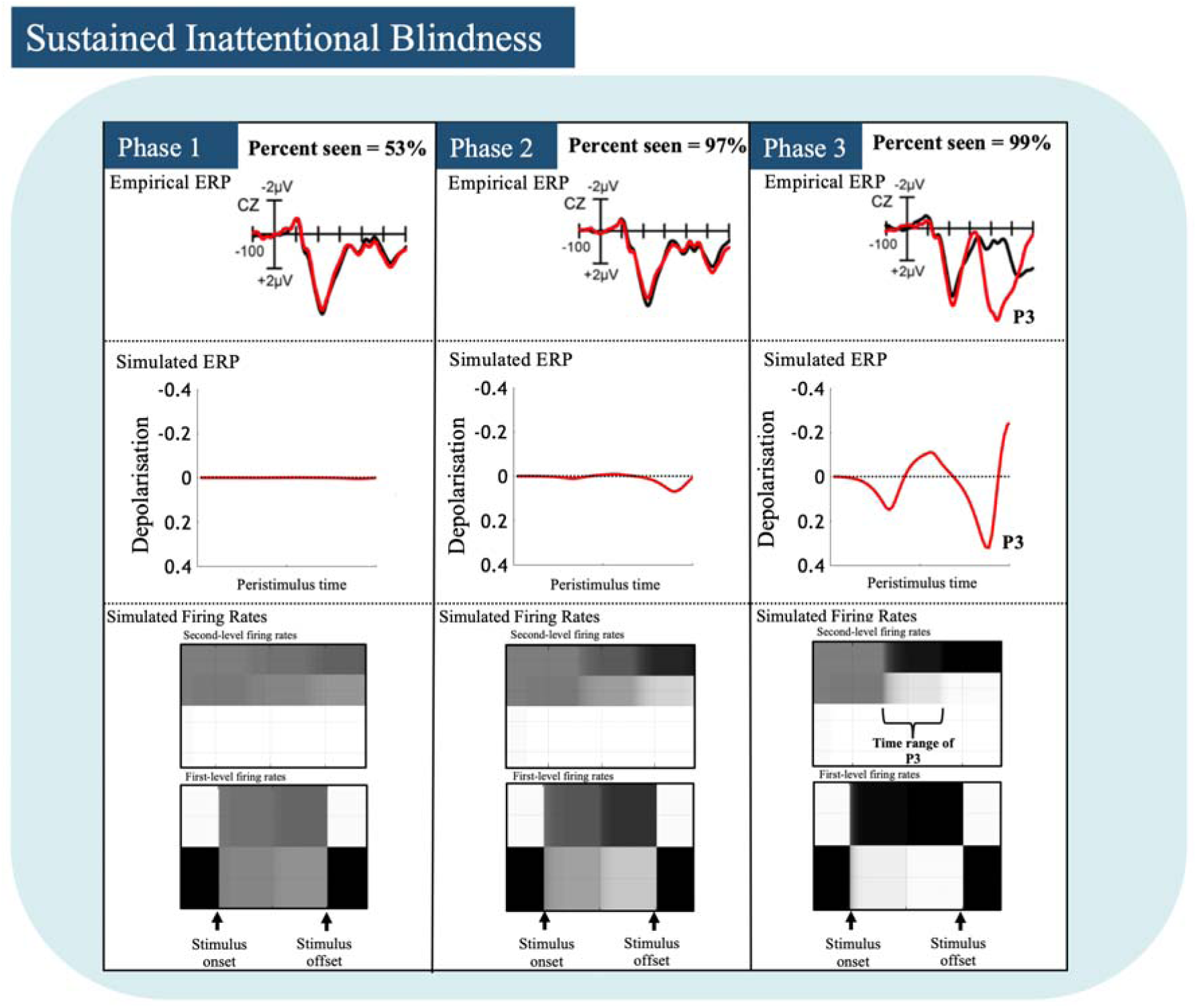
Report frequency, simulated ERPs, and associated firing rates predicted for each of the three phases of the sustained inattentional blindness task. The empirical ERP plots, taken from the original study conducted by Pitts et al (2014), show the amplitude for “random bar” (black) versus “square present” (red) trials at electrode CZ. Firing rate plots illustrate how more gradual updates in 2nd-level beliefs do not produce P3-like ERPs (middle panel, Phase 2) despite self-reported experience of the square, while a faster rate of change in 2nd-level beliefs does produce P3-like ERPs (right panel, Phase 3) in addition to almost identical self-reported experience.

This result is noteworthy. The dissociation of the P3 and visual consciousness emerges naturally out of the belief updating scheme that underwrites the PGNW. Further, it leads to a straightforward prediction; visual consciousness will be accompanied by a late ERP whenever confidence in a particular state at a high level of the hierarchy changes rapidly.

Having shown that our model reproduces the minimal contrast results cited in support of the original formulation of the GNW, explains away otherwise contradictory results in the minimal contrast literature, and accounts for the dissociation of the P3 and visual awareness, we now turn to the role of visual expectation – and illustrate how our model offers an extension of the previously introduced four-way taxonomy of factors underlying conscious report. In addition to attention and stimulus strength, our model introduces a third factor: trial-by-trial changes in prior visual expectation.

### 3.4 Extended Taxonomy: Expectation and Visual Consciousness

There is now a large body of evidence showing that expectation plays a fundamental role in determining the content of visual consciousness. Specifically, under conditions of continuous flash suppression predictive cues accelerate the entry of a suppressed stimulus into consciousness (van Gaal et al., 2015). In binocular rivalry paradigms, predictive context increases the dominance of stimuli congruent with that context (Denison et al., 2011; Valuch & Kulke, 2019). Cross-modal predictions also accelerate the re-entry of stimuli into consciousness after a period of motion induced blindness (Chang et al., 2015). In the absence of attention, expectations reliably induce illusory perception of absent stimuli (Aru et al., 2018). And, when viewing ambiguous figures, expectations have been shown to bias the perceived direction of rotation (Sterzer et al., 2008). In light of this, if visual consciousness is to be fully understood, it appears essential to extend the taxonomy of factors underlying conscious access to include expectation.

To integrate the role of expectation – and the violation of expectation – into the taxonomy, we altered the prior probability of the “square present” sequence in the second level **D** vector for each of the four parameter settings used in the four-way taxonomy, such that the model was approximately twice as confident, a priori, in either the “square present” sequence (“consistent prior expectation” condition) or the “random bar” sequence (“inconsistent prior expectation” condition). We treated each trial as independent, so the manipulation of prior expectations most plausibly corresponds to the use of explicit cues.

We retained the same generative model architecture used in the previous two sections, allowing us to independently manipulate expectation, attention, and stimulus strength. We are aware, however, that independently manipulating these factors in an experimental setting is far from trivial. In the interest of making our model empirically useful, we end this section by proposing a novel Posner cueing paradigm aimed at empirically validating the results of our simulations.

Finally, it must be highlighted that the behavioural results of the following simulations should be interpreted as directional hypotheses, as opposed to precise predictions about the percentage of seen vs unseen trials. In contrast to firing rates and ERPs, the model’s report behaviour (policy selection) depends on a number of parameters (policy precision and motor stochasticity) that need to be fitted to each participant individually and will substantially shift the size of the effect for manipulations of prior expectation.

#### 3.4.1 Extend Taxonomy: Simulated Behaviour

We presented the model with 800 “square present” trials, 100 corresponding to “consistent prior” and “inconsistent prior” conditions, for each of the four parameter settings used in the four-way taxonomy. Across all taxonomy conditions, we found that consistent prior expectations increased both the accuracy of forced-choice behaviour and the percentage of trials where the model reported having seen the stimulus (see figure 7 for percentages). Similarly, inconsistent prior expectations decreased both the accuracy and the number of seen trials. The effects were particularly pronounced in all of the otherwise sub-threshold conditions. This result makes intuitive sense, as expectations will have the greatest enhancing effect on visibility when a stimulus is marginally below the threshold for “ignition”; and, in the absence of attention and/or precise sensory input, top-down messages will dominate perception.

**Figure 7.**
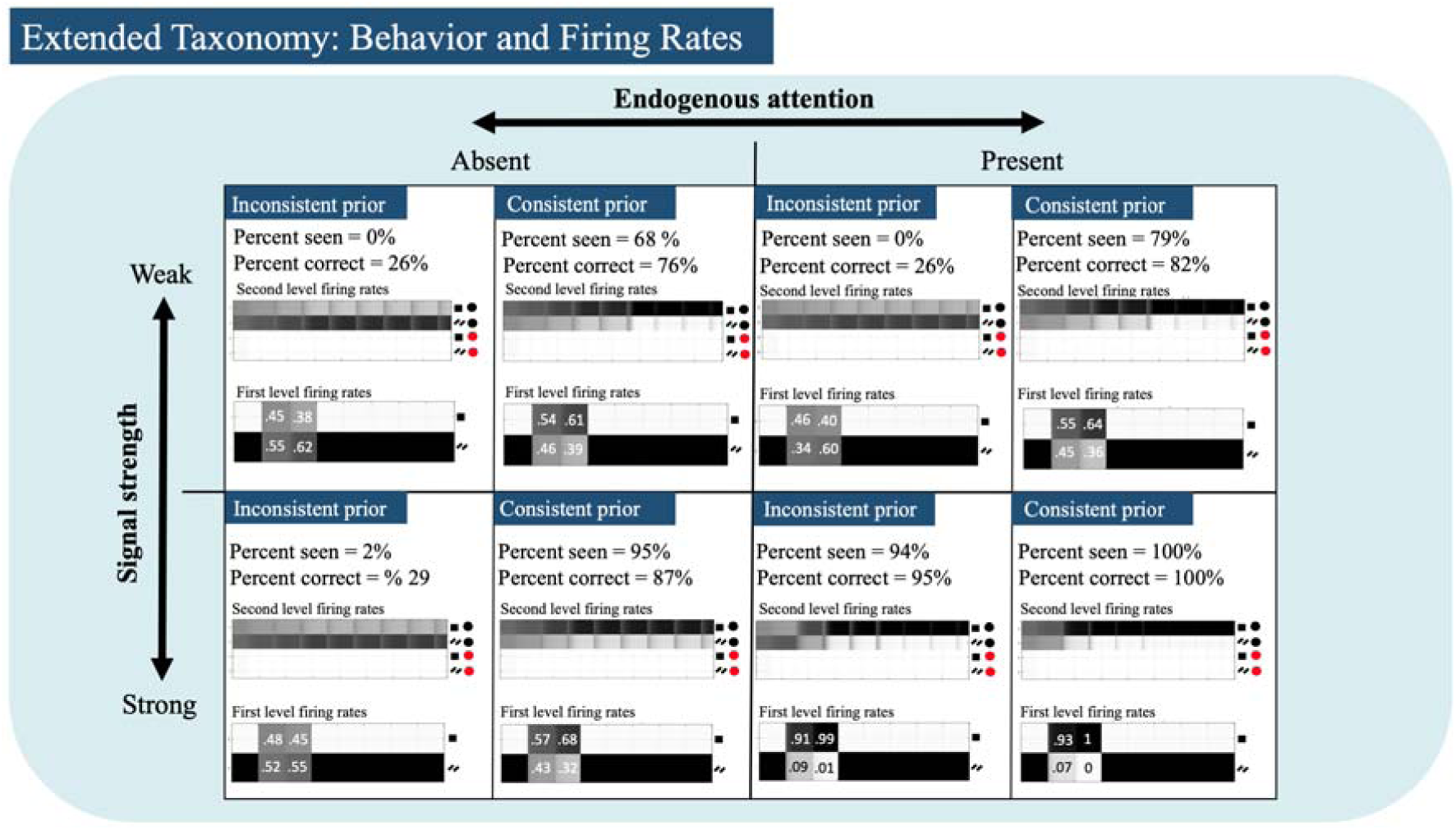
Report frequency, forced choice behaviour, and simulated firing rates predicted for each consistent-inconsistent prior combination of the quadrants shown in the four-way taxonomy. Relative to the results of the four-way taxonomy, consistent expectations increased forced choice accuracy, the percentage of trials reported as “seen”, and boosted the enhancing effect of feedback from the higher level. Inconsistent priors had the opposite effect, reducing accuracy, the percentage of “seen” trials and first level firing rates.

While these results are consistent with a number of studies showing an enhancing effect of consistent prior expectations for both forced-choice performance and stimulus detection (Aru et al., 2016; Stein, & Peelen, 2015), there is some evidence showing that when expectations are induced by explicit cues, they boost subjective visibility but do not alter accuracy (Andersen et al., 2019). Similarly, Lamy et al (2017) found that prior experience of a target increased visibility but did not alter response priming. Here forced-choice behaviour, like subjective report, depends on policy selection at the second level of the model. However, if forced choice behaviour depends upon a distinct neural substrate that operates over a shorter timescale than subjective report, it would be better modelled by policy selection at the first level of the model (c.f. Maniscalco & Lau, 2016). If this were the case increasing the prior probability of a state at the second level (as we have done here) would have a marginal effect on forced-choice behaviour. This represents an important possible extension of our model architecture that will be addressed in future work.

#### 3.4.2 Extending the Taxonomy: Simulated Firing Rates

Next, we simulated the firing rates associated with consistent and inconsistent priors. In all sections of the extended-taxonomy, relative to the four-way taxonomy, consistent priors enhanced the amplifying effect of feedback. Correspondingly, inconsistent priors dampened the effect of feedback. Interestingly, in the “strong signal + attention present” condition, although inconsistent priors dampened the amplifying effect of feedback (i.e., in comparison to the four-way taxonomy and the consistent prior condition), the net effect of feedback still raised the firing rate of first level neuronal populations.

Again, this makes intuitive sense; feedback is driven by posterior confidence at the second level, and, in all but the “strong signal + attention present” condition, the sensory likelihood mapping is relatively imprecise. As such, posterior confidence at the second level is dominated by the effect of prior expectation. Consistent expectations increase second-level posterior confidence to a sufficiently large degree to shift otherwise unconscious trials over the threshold for “ignition” by magnifying the feedback to the first level (while inconsistent priors reduce posterior confidence and dampen top-down feedback). In contrast, when the first-level likelihood mapping is more precise, as is the case in the “strong signal + attention present” condition, inconsistent priors carry less influence and second-level posterior confidence in the “square-present” sequence is still high enough to enhance top-down feedback (i.e., despite it being dampened relative to the four-way taxonomy and consistent prior condition).

This result produces three novel predictions. First, when consistent expectations raise posterior confidence past the threshold for ignition, feedback from frontoparietal to sensory regions will be enhanced. Second, when a stimulus is well above threshold, inconsistent expectations will reduce the amplification of feedback relative to consistent expectations. Third, based on the extrinsic connectivity predicted by the neural process theory (see figure 1), feedback from frontoparietal to sensory regions will be associated with a specific pattern of effective connectivity.

That is, consistent prior expectations are predicted to disinhibit granular layers in the relevant neuronal populations in sensory cortices (via feedback connections originating in superficial pyramidal cells in frontoparietal regions), while inconsistent priors are predicted to inhibit granular layers in the same lower-level neuronal populations. This last prediction, although highly specific, can be readily tested via dynamic causal modelling (e.g. Parr et al., 2019b). However, it is important to note that this prediction is dependent upon the use of variational message passing (or marginal message passing), and other message passing schemes have been proposed (see Parr et al., 2019a).

#### 3.4.3 Extend Taxonomy: Simulated Event Related Potentials

Lastly, we simulated the ERPs predicted by our model for all combinations of consistent and inconsistent prior expectation conditions within the four-way taxonomy (see figure 8). Despite the previously described changes in reported visibility, in most cases ERPs were relatively unaffected by consistent and inconsistent priors at both levels of the model. However, in the “strong signal + attention present” condition the amplitude of ERPs at the second level of the model, corresponding to the P3, were dampened by a consistent prior and boosted by an inconsistent prior relative to neutral priors. This result makes sense, because in the consistent prior condition the model was already confident that it would be presented with a square, so the presentation of the stimulus caused a smaller and less rapid belief update, while the opposite was true in the inconsistent prior condition – thus reducing the rate of change of posterior expectations at the second level in the consistent prior condition and increasing it in the inconsistent prior condition.

**Figure 8.**
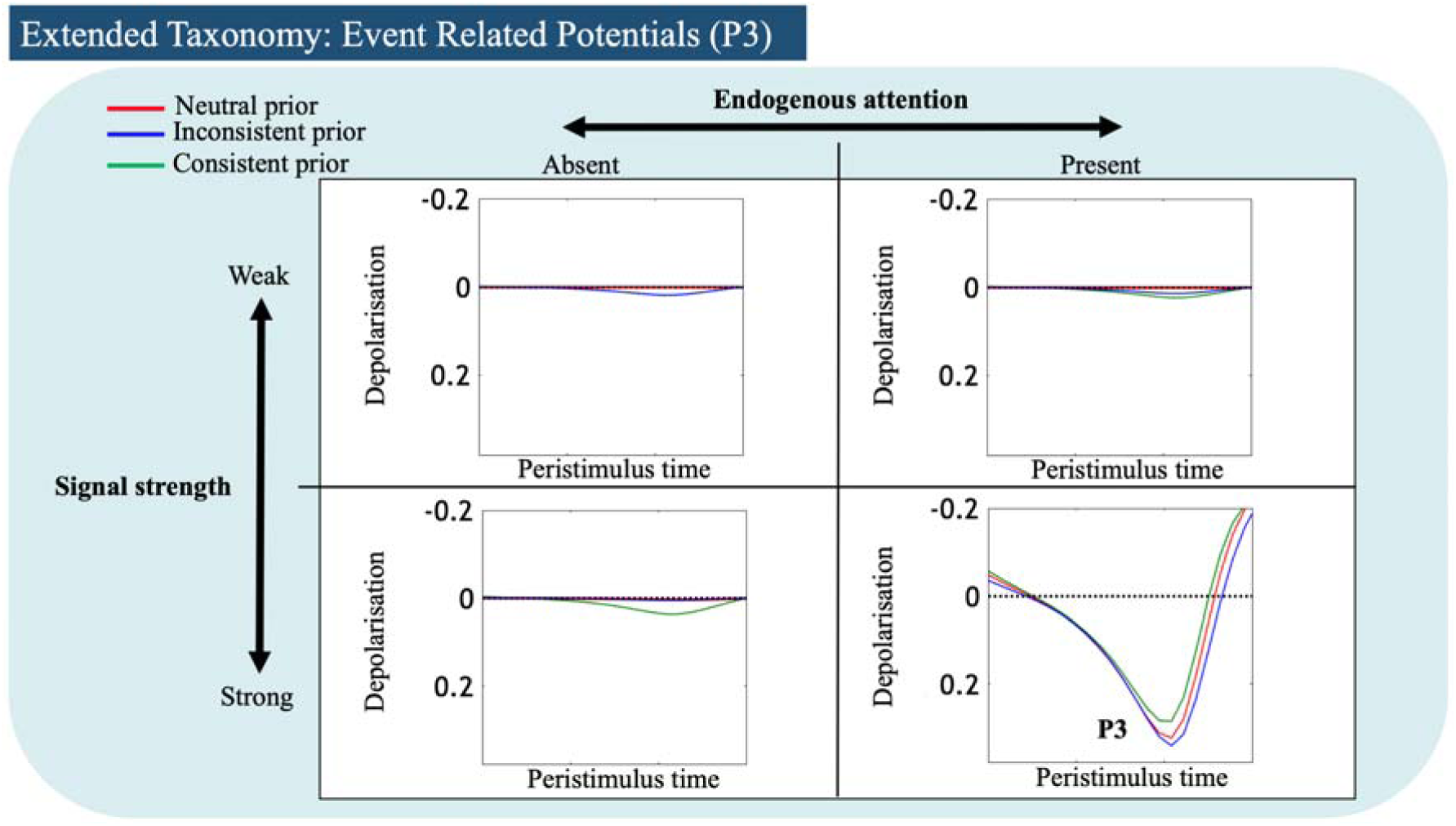
Second-level ERPs predicted for each combination of consistent or inconsistent priors with each quadrant in the four-way taxonomy. Notice that 1) the P3 is attention- and stimulus strength-dependent (i.e., only the bottom-right quadrant shows clear responses above noise levels), 2) it is enhanced by inconsistent priors, and 3) it is dampened by consistent priors.

Importantly, these results lead to another novel prediction. Specifically, contingent upon the presence of attention, and sensory input being sufficiently precise, a consistent prior should decrease the amplitude of the P3 (relative to neutral and inconsistent prior conditions), whilst an inconsistent prior should increase the amplitude of the P3 (relative to neutral and consistent prior conditions). Indeed, this prediction has already been partially confirmed empirically. Melloni et al. (2011) found that expectations induced via a history of prior exposure to a stimulus both increased the proportion of trials reported as seen and decreased the amplitude of the P3. However, this study did not simultaneously manipulate attention, which will be a critical further test of the hypothesis.

#### 3.4.4 Extend Taxonomy: A Novel Paradigm for Dissecting the Influences of Signal Strength, Attention, and Expectation on Conscious Access

The structure of our generative model is generic enough to generalise across paradigms that involve inference; however, as noted above, in an empirical setting independently manipulating expectation, signal strength, and attention poses a number of methodological challenges, with expectation often being confounded with attention (e.g. Rahnev et al., 2011). In the interest of making the predictions of our model as straightforward to test as possible, we here outline a possible extension of the Posner cueing paradigm introduced by Kok and colleagues (2012) that would allow for the independent manipulation of expectation, signal strength, and attention.

The key feature of the design is the orthogonal manipulation of expectation and attention (see figure 9). Expectation can be manipulated in a block-wise manner with a predictive cue appearing at the beginning of every block consisting of a word (“left”, “right” or “neutral”) indicating the likelihood with which the stimulus will appear in a particular hemifield on each trial. Attention, in contrast, will be manipulated in a trial-wise manner, with a cue appearing at the start of every trial indicating the hemifield to which the subject should covertly direct their attention. However, the attention cue will contain no information about the likelihood of the stimulus’ location. Finally, stimulus strength will be manipulated by altering the time between the stimulus and the backward mask. Each block would begin with a predictive cue, and each trial would begin with the presentation of an attention cue, followed by a briefly presented stimulus (a grating in the above figure) that is congruent with the prediction on 75% of trials (paired with either a backwards mask or a blank). Subsequent to the presentation of the mask (or blank), subjects would be given a forced-choice task and asked to provide a subjective report.

**Figure 9.**
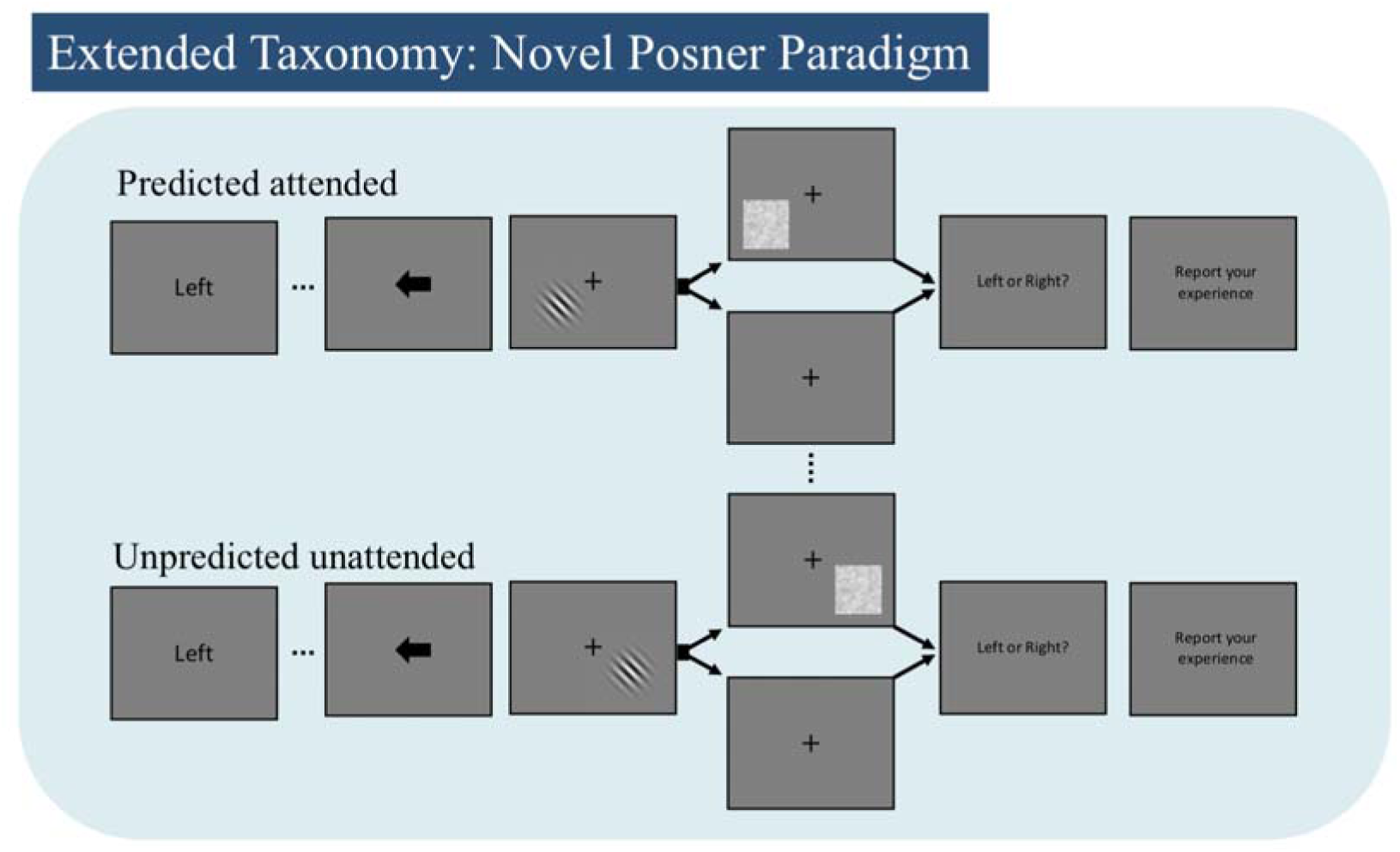
A potential extension of Posner cueing paradigm introduced by Kok and colleagues (2012) that could allow for the independent manipulation of expectation, stimulus strength, and attention. By manipulating expectation in a block-wise manner, and attention and stimulus strength in a trial-wise manner, the paradigm would allow all twelve combinations of expectation (consistent, neutral, inconsistent) by attention (present, absent) by stimulus strength (strong, weak) to be studied within one paradigm. Shown above are predicted/attended and unpredicted/unattended combinations.

Crucially, since expectation, attention, and stimulus strength are manipulated independently of each other, this paradigm could allow all 12 quadrants of the extended taxonomy to be studied within one paradigm.

## 4. Discussion

### 4.1 The role of deep temporal structure

The defining (measurable) feature of conscious access is subjective report (Baars, 1988; Dehaene, 2014; Fleming, 2019), which requires the coordination of processing between perceptual, motor, and auditory-verbal systems, all of which evolve over a rapid temporal scale. The core notion underlying our model is that conscious access is a fundamentally inferential process that can *only* occur at a level of processing that is sufficiently temporally deep to integrate information from lower levels of the hierarchy and contextualise processing at these lower levels. To produce subjective reports, a system must infer the state of a lower level perceptual system, integrate this information into a representation that is not tethered to the moment by moment sensory flux, and use this representation to modulate more controlled, slowly evolving trajectories of action over time. Put another way, temporally deeper levels are necessary to encode patterns of covariance in lower-level sensory and motor representations over time under different goal states. Without a sensory representation updating predicted patterns of covariance at this higher level (to a sufficient degree), the use of that piece of sensory information by the more complex cognitive processes carried out at the higher level would be greatly limited (i.e., only promoting implicit biases through small changes in higher-level posterior distributions). Technically, the insight here is that ignition and the global workspace are descriptions of evidence accumulation or assimilation – which necessarily implies some temporal narrative. The nature of this narrative turns a straightforward Markovian model of the world into a semi-Markovian model with deep temporal structure.

The first major insight afforded by our model is that many previous electrophysiological results can be reproduced based only on assuming a simple 2-level model with deep temporal structure. Self-reported conscious vs. unconscious percepts can then be accounted for by specifying the integrative functions of the higher level of the model that are plausible based on the known neural correlates of consciousness, and how the different hidden state factors (which could perhaps correspond to distributed processing hubs within association cortices implicated in domain-general cognition i.e. van den Heuvel et al., 2012) operate on those contents. The second major insight comes from how our model illustrates the way prior expectation can fit seamlessly within this structure – affording a number of novel, testable predictions.

### 4.2 Relationship to Other Models

The PGNW is a formal extension of the GNW and, as such, the models share many similarities. However, as should be clear by this point, the PGNW diverges substantially from the original GNW model. Ultimately, the point of departure for the PGNW is its implementation in an Active Inference architecture, which, as we have shown, has a number of important consequences. Foremost, by leveraging the process theory that accompanies Active Inference, we are able to reproduce/explain previous findings and make predictions about the neurobiological implementation of the inferential machinery that we argue underlies conscious access. Crucially, this also allows the PGNW to make specific predictions about the role of visual expectation. In contrast, the inferential machinery cited in support of the standard GNW model (i.e. King & Dehaene 2014) remains at a more abstract computational level of description (in the sense of Marr, 1980). What the PGNW retains is the fundamental insight that conscious access makes information widely available to domain-general cognitive processes (i.e., represented by the different state factors at the second level of our model). It is for this reason that we retain the workspace label. However, unlike the initial conceptual account of the PGNW introduced by Hohwy (2013) and Whyte (2019) which, as Marvan and Tomáš (2020) point out, relies on explanatory machinery external to the Active Inference framework to explain conscious access, here we identify the global availability of information with temporally deep processing, and conscious access with the posterior confidence threshold required for report (broadly construed as goal-directed verbal report, button presses, saccades or any other method of goal-directed reporting of subjective content). As such the PGNW explains conscious access exclusively in terms of the explanatory tools of Active Inference.

The model that is most similar to the PGNW is Fleming’s (2019) higher-order state-space (HOSS). Both the PGNW and HOSS are implemented in hierarchical generative models, and as such they entail similar predictions. However, there are two key points of separation. First, HOSS casts conscious access as a metacognitive inference about the presence or absence of a stimulus in the content of a generative model of perception. This inference relies on an abstract state representing presence or absence that is independent of the specific content of a stimulus. The function of this metacognitive state is to differentiate stimulus representations from noise distributions at lower levels of the model (c.f. Lau, 2007, 2019). A stimulus representation becomes available for report according to HOSS when the model infers that a distribution is different enough from a noise distribution to be classified as present.

There is a sense in which HOSS is simply a higher-order version of the PGNW. In fact, it would be relatively simple to introduce a new hidden state factor corresponding to the presence or absence of a stimulus and factorise a generative model such that the presence of the stimulus would be independent of the content of the stimulus. Computationally, however, this state would have no function. Precision estimation is an inbuilt feature of Active Inference architectures (see Parr & Friston 2017 for technical details) that modulates the updating process in response to the estimated reliability of a bottom-up signal, without needing to posit an additional abstract state representing presence and absence.

This brings us to a second key difference. According to the HOSS model, absence of a stimulus is explicitly represented in addition to all the possible states of a stimulus making the state space asymmetric. Fleming (2020) leverages this asymmetry to explain the ignition response that sweeps across frontoparietal cortices during conscious access. Because there are many more ways that a stimulus can be present than absent, the presence of a stimulus causes larger belief updates than when a stimulus is absent. Although this is ultimately an empirical question, we regard the explicit representation of the absence of a stimulus in the perceptual state space to be a somewhat unrealistic idealisation. Participants in minimal contrast paradigms are fully aware of the presence of the background screen (itself a stimulus) and the task requirements, and frontoparietal cortices are, of course, active the whole time. The wide scale activation of these regions when a stimulus is seen is only apparent because we isolate the processes underlying the reportability of the stimulus, while holding all of these other variables constant. When participants do not report a stimulus, they are not perceiving “absence”; at a minimum the content of perception will include the background screen. It may be a mistake, therefore, to explicitly represent the absence of a stimulus in a generative model of perception. Instead, as we have done in the current model, the state space should consist of sequences of stimuli with and without the critical stimulus. The ignition response is then instead explained by the update that occurs at temporally deep levels of the model when the stimulus is seen as opposed to unseen.

Finally, it is worth mentioning that, although there have been proposals aimed at integrating other major theories of consciousness, such as integrated information theory (Oizumi, Albantakis & Tononi, 2014) with VFE and active inference (Safron, 2020a, 2020b), these proposals remain largely at the level of conceptual analysis and do not afford the detailed simulations and resulting empirical predictions that are a straightforward consequence of the PGNW architecture. However, it remains an open question whether or not systematic relationships might be found between measures of integrated information and VFE in future work (for discussion on this topic see Friston, Wiese & Hobson, 2020).

### 4.3 Brief Note on Phenomenology and the Phenomenal Consciousness - Access Consciousness Distinction

Although not the focus of this paper, it is worth briefly clarifying how phenomenology does and does not plausibly situate within our model. Specifically, we would like to avoid implicitly conveying that the phenomenological contents of our first-person experience correspond to the contents of second-level states. A major problem with this is that the contents of second-level states appear to operate on timescales that are too slow to match the moment-to-moment sensory flux of perceptual experience.

However, there is also a problem with identifying phenomenological contents with the contents of lower-level states. To see why, consider that, as is the case in our model, there is no explicit representation of absence at the higher level of the model that generates reports. The model will always represent and report an experience of something based on the posterior distribution over second-level states (e.g., either lines or a square; and note that the model’s state space could also be extended such that the agent could report the experience of ‘just seeing the background screen’). If this architecture is representative of human cognition, it highlights an interesting change in perspective. Specifically, the question about phenomenology being separable from access (c.f. Block, 2005) changes to a question about the possibility of inconsistencies between phenomenology and *what* was accessed (i.e., the states represented at the second level). For example, if the agent reported currently seeing lines (and not a square), and yet a square stimulus was present and encoded at the first level, a strong distinction between phenomenology and access would not merely entail that “square” phenomenology was present but not accessed. Instead it would mean that the agent’s confident self-reported phenomenology of experiencing lines (i.e., what was represented at the second level) would be inconsistent with their “true” phenomenology of experiencing a square (i.e., based on what was represented at the first level). In other words, they would be wrong about what they believed they were currently experiencing or had just experienced. Taken to the extreme, this would entail that any honestly reported phenomenology could be problematically different from true phenomenology (e.g., a person honestly reporting experiencing a loud screeching sound could have the “true” phenomenology of a hearing a piece of classical music).

Another way to highlight this problem more formally is by considering that one could manipulate the second-level likelihood mapping (**A** matrix) in our model while leaving the first-level likelihood mapping unchanged (i.e., one could selectively manipulate the nature of the messages that are passed from the first level to the second level that update higher-level beliefs at each timestep). If so, first-level states would still reliably track presented stimuli (e.g., square stimuli would activate first-level square states), but those states could now update the second-level in an entirely different way. For example, with the right second-level likelihood mapping, red circle and line representations at the first level could be specified so as to pass messages to the second level that activate representations of, and promote self-reported phenomenology of, a black circle and a square (which would also obviously be problematic for the empirical study of conscious experience; i.e., the presence of a particular phenomenology would become empirically unfalsifiable).

To avoid this uncomfortable conclusion, while also keeping sensory phenomenology at the correct timescale, we suggest that phenomenology in our model is most plausibly situated at the point during which (and based on the manner in which) lower-level representations *update* the content of higher-level states. In other words, phenomenology in our model most plausibly depends on the nature of the messages passed from the lower to the higher level, and would occur at the point where the higher level *assesses* or “decodes” the contents of the lower-level signals through the second-level likelihood mapping (**A** matrix).

These updates occur at the fast timescales associated with the sensory flux; yet, the nature of each fast-timescale update (i.e., the nature of the influence the first level has on the second level at each time point) will necessarily be correlated with self-reported beliefs about what type of phenomenology was experienced – preventing the possibility of strong disagreements between “true” and self-reported beliefs about phenomenology (for more on this line of argument see Smith, 2016). Note that the focus on belief updating thus speaks to phenomenal consciousness as a process of inference.

Thus, the perspective motivated by our model might therefore be seen as somewhat at odds with a strong distinction (or at least with the most commonly made distinction; (Block, 2005)) between phenomenal consciousness and access consciousness. In contrast to the typical distinction, our model suggests that (empirically verifiable) phenomenology and access consciousness each rely on particular (partially theoretically separable) types of access (c.f. Cohen & Dennett, 2011). Phenomenology (i.e., the content of first-person experience) most plausibly depends on the faster timescale process by which first-level representations update higher-level beliefs via the nature of the messages passed through the second-level likelihood function (**A**-matrix). In contrast, access consciousness (as typically defined), and all of the functional benefits that it is associated with, corresponds best to the encoding of posterior beliefs over hidden states at the second level, which, although updated by the moment-to-moment sensory flux, are themselves representations of regularities that occur over longer timescales.

A final point worth emphasizing is that, while verbal or other types of goal-directed self-report measures remain the gold-standard in assessing the presence or absence of conscious experience, our model also clearly distinguishes access consciousness from report. That is, access consciousness in our model depends only on the precision of posterior beliefs at the higher level. In a no-report paradigm, for example, an active inference task model analogous to the one we have presented would be able to demonstrate that – even if no report was made at the timepoint of stimulus presentation – posterior beliefs were sufficiently precise at that timepoint such that, *had the person been incentivized to report their experience*, they *would have* reported seeing the stimulus. Such extensions of our model to no-report paradigms represents an important future research direction.

### 4.4 Limitations

To make the PGNW testable, we have deliberately limited the scope of the model to experimental settings where visual consciousness is operationalised via report. We follow Baars (1988) in taking report, or rather the availability of information for report, as the epistemic foundation of the scientific study of consciousness. However, we acknowledge that report paradigms come with methodological difficulties (Tsuchiya et al., 2015). By the same token, in limiting the scope of the PGNW to the visual modality, we are aware that we are reifying the pervasive bias in consciousness science of primarily studying vision. That said, it is crucial to emphasise that the model structure is sufficiently general that it can be straightforwardly applied to other modalities (e.g., first-level observations could fairly easily be understood as auditory as opposed to visual). A somewhat similar 2-level architecture was also recently employed to simulate emotional awareness based on interoceptive stimuli (Smith, et al., 2019a).

The main focus of the GNW has, until recently, been the contents of consciousness (especially vision). Likewise, this paper has limited the scope of the simulations to conscious access in awake behaving subjects. However, there is now a growing body of work showing that the GNW is also able to account for differences in levels of consciousness (see Mashour et al, 2020 for a review), in the sense that the integrity of connectivity between workspace nodes covaries with changes in the overall state of consciousness. Specifically, all major classes of general anaesthetic have been shown to in some way disrupt frontoparietal networks (Hudetz & Mashour, 2016). Further, work in non-human primates has found that, only in a waking state, GNW nodes – including parietal, prefrontal, and sensory cortices – display a wide range of functional connectivity patterns much richer than anatomical connectivity. In contrast, a range of anaesthetics with different molecular mechanisms (ketamine, propofol, and sevoflurane) all drastically limit functional connectivity to patterns that resemble anatomical connectivity (Uhrig et a, 2018). The simulations in this paper do not speak directly to these results, although they do sit well with the relevant aspects of the computational architecture of the PGNW. For example, the most plausible analogue in our model to the disconnection between prefrontal and parietal cortices that accompanies general anaesthesia would be a lesion to the 2nd level A-matrix (i.e., such that the first and second levels no longer exchange information). In this case, second-level representations would quickly become maximally imprecise (barring infinitely precise second-level priors) and carry no information, plausibly corresponding to general unconsciousness (i.e., no conscious content of any kind). Importantly, the agent would also become insensitive to longer timescale regularities in sensory input, explaining the abolishment of the P3 response to violations of longer time scale auditory regularities during sedation and sleep (Shirazibeheshti et al, 2018; Strauss et al, 2015).

In addition to these big picture limitations, our model has a number of more specific limitations that apply strictly to the study of visual consciousness. Principally, our model only has two levels and we treat the entire visual system as a singular and discrete level instead of modelling it for what it is – a continuous and multi-level system. The need for multiple levels brings us to the next limitation. As Kouider et al (2010) argue, people are often only partially aware of a visual scene in the sense that they may be aware of an object’s colour but not its identity. This requires information to skip levels of the hierarchy, which is also not possible in the present model. A more complete model would therefore allow both shallower and deeper representations to selectively update the second “workspace” level (e.g., allowing separable awareness of representations of an eye vs. a face vs. a person’s identity).

## 5. Conclusions and Future Directions

This paper introduced a formal extension of the global neuronal workspace – the predictive global neuronal workspace – implemented within a deep Active Inference architecture. In addition to explaining and unifying otherwise disparate findings in the neural correlates of visual consciousness literature, the predictive global neuronal workspace model presented here generates several empirical predictions, and mechanistic neuro-computational explanations, regarding the relationship of the P3 and subjective report, the neurobiological implementation of the inferential machinery underlying conscious access, and the role of expectation in visual consciousness.

In future work, we hope to build on the wealth of existing Active Inference models (e.g. Allen et al., 2019; Parr et al., 2019c; Smith et al., 2019a; 2019b), to extend the PGNW to other sensory modalities and more sophisticated experimental paradigms.

### Software note

The generative model detailed in this paper used a generic belief updating scheme (**spm_MDP_VB_X**.**m**) implemented in Matlab code using the freely available SPM academic software: https://www.fil.ion.ucl.ac.uk/spm/. The scripts used to produce the specific simulations reported here can be downloaded from https://github.com/CJWhyte/PGNW_ERP-1_2020.

## Conflict of interest statement

The authors have no conflicts of interest to disclose.

## Appendix Full Model Specification

At the first level of the model (see figure A1), the **D** vectors specified the initial state of the four hidden state factors; top-down attention (present, absent), internal stimulus (bars/square), peripheral stimulus (red/black circles), and auditory-verbal states (single words: “silent”, “I”, “see”, “a”, “square”, “didn’t”, “anything”). The state transitions specified by the **B** matrices were all identity matrices, meaning that the hidden states were stable across the course of each trial. The likelihood mapping between the hidden states and outcomes, specified by the **A** matrices, is where we implemented the attention and signal strength manipulations. The peripheral stimulus and language matrices were both fully precise (identity matrices). In contrast, we reduced the precision (denoted by (for stimulus strength and Σ for attention) of the mapping between the internal state and the outcomes by passing what were initially identity matrices through two softmax functions controlled by precision parameters representing the effects attention and signal strength (i.e. presentation time). Higher values of these parameters made the **A** matrices more precise. We set up the interaction between the **A** matrices such that the likelihood mapping for the internal stimulus factor was more precise when the agent was in an attentive state. In contrast, stimulus strength manipulations reduced the precision of the mapping between stimuli and hidden states independent of attentional state.

**Figure A1.**
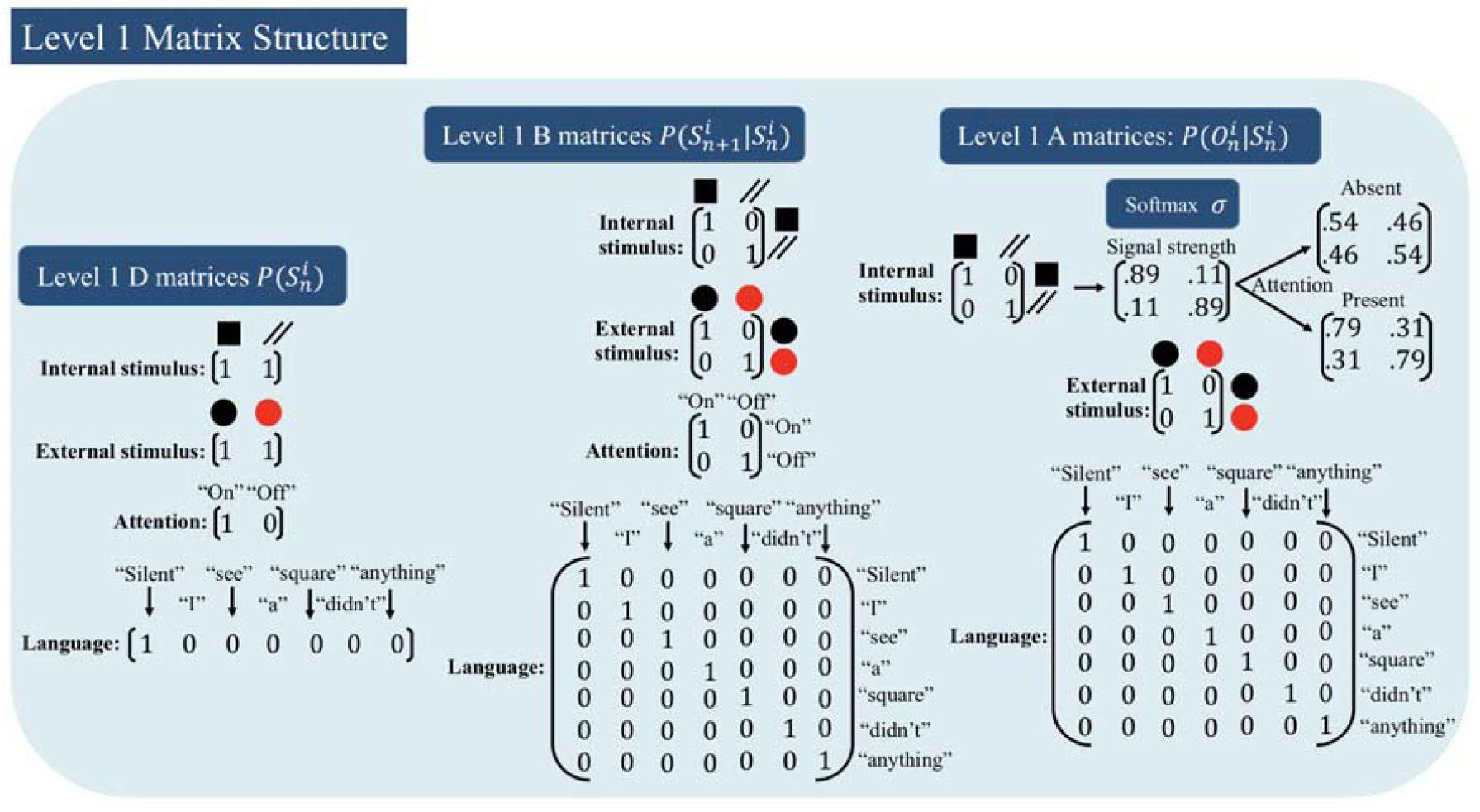
1st-level generative model matrices. All matrices are passed through softmax functions such that the columns of each matrix, and the rows of each vector, always sum to one. Each column of the **D** vector represents the probability of a hidden state. Columns of the **B** matrix correspond to states at time *t* and rows correspond to states at *t+1*. Here all the **B** matrices are identity matrices meaning that the states were believed to be stable across each trial. Columns of the **A** matrix correspond to hidden states while rows correspond to observations. An identity mapping therefore implies a deterministic (i.e. precise) likelihood mapping between states and observations. To model the effects of attention and stimulus strength the **A** matrix encoding the likelihood mapping for the internal segment of the stimulus was passed through a softmax function twice with a precision multiplier representing attention () which could be present or absent, and stimulus strength which could be strong or weak (. The combined effect of the attention and signal strength multipliers determine the final precision of the matrix as depicted above. With the exception of the **A** matrix encoding the likelihood mapping for the internal segment of the stimulus, all other first-level **A** matrices were identity matrices, meaning that the mapping was deterministic. Finally, it is important to note that, for visual simplicity, the matrices displayed above have not been factorised (in the code, these matrices are Kronecker tensor products).

At the second level (see figure 5) the three hidden state factors specified by the **D** vectors were: sequence type (black disk and square, black disk and bars, red disk and square, red disk and bars), time point within trial (1-8), and report state (wait, seen, unseen). We set the initial level of the report state to “wait”. The **B** matrix for sequence type was an identity matrix, meaning that the agent believed a priori that the sequence type would not change mid-trial. Trial phase was set up such that time point 1 transitioned to time point 2, which transitioned to time point 3 and so on until the end of the trial. For time points 1-4, all states in the “report” **B** matrix mapped to “wait”; however, at time point 5 the agent had control over the **B** matrices for the report state, meaning that it could transition to either a “seen” or “unseen” state depending on which policy best minimised expected free energy. The **A** matrices were factorised such that the mapping from hidden states to outcomes was dependent on the time point in the trial (see the time-in-trial hidden state factor in figure A2). In other words, the narrative of this paradigm was specified in terms of interactions between time and other content-specific hidden state factors (i.e., and the interaction between ‘when’ and ‘what’). For example, at time point 1 both square sequence and bar sequence hidden states predicted a bar outcome (recall that second-level outcomes are also first-level hidden states). While at the 2nd and 3rd time points the square sequence predicted a square outcome and a bar hidden state predicted a bar outcome. At time point 4, both the square sequence and bar sequence once again predicted a bar outcome. To model the recurrent feedback between hierarchical levels characteristic of ignition, the square sequences mapped to the square outcomes for time point 2 and 3 – allowing the state at the second level to influence belief updating at the first level via the second-level **A** matrix. From time points 1-4 all the states in the “report” factor mapped to the “silent” first-level verbal state. However, from time point 5 on, “seen” and “unseen” states entailed a different sequence of lower-level word representations. The “seen” report state entailed the words (in order) “I” “see” “a” “square” at each successive time point, while the “unseen” report state entailed the words “I” “didn’t” “see” “anything” in that order. The report state also had a likelihood mapping to a feedback outcome. The agent was “correct” if, at time point 8, they reported “seen” after a square sequence or “unseen” after a bar sequence, and “incorrect” if they reported “unseen” after a square sequence or “seen” after a bar sequence (this preferred feedback was used to motivate honest verbal reporting policies; see below). To account for the diffuse nature of feedback projections (Bannister, 2005; Garcia-Cabezas et al., 2019), we lowered the precision of the **A** matrix for the “sequence type” factor (precision = 0.8) providing a plausible threshold on ignition events.

**Figure A2.**
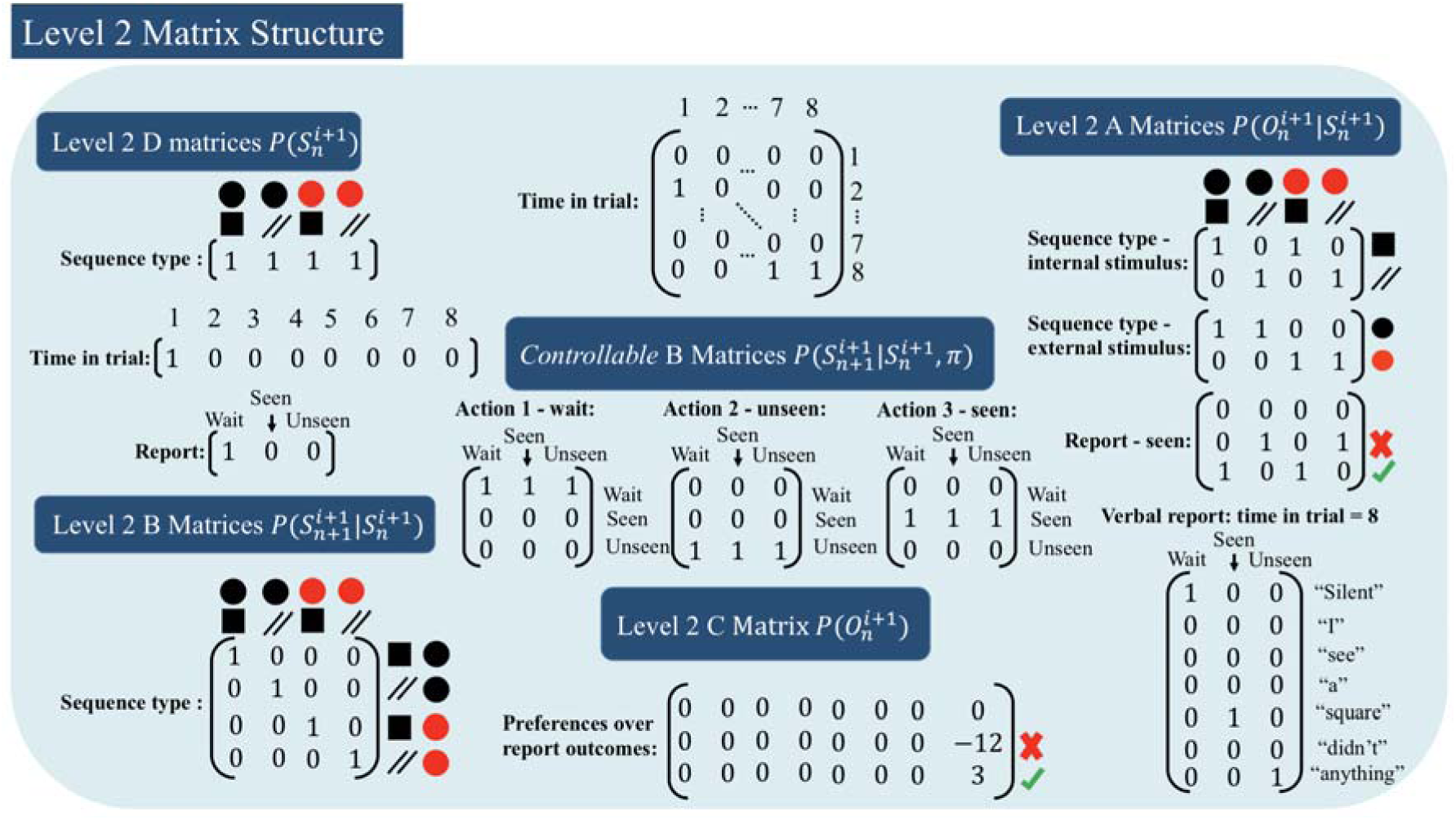
2nd-level generative model matrices. **D** vectors for sequence type (unless otherwise indicated) assigned equal probability to each state. The **D** vector for the trial phase hidden state factor was initialised so that the model would always start each trial with full confidence that it was in the state corresponding to time point 1. Similarly, the **D** vector for the report state was initialised so that the model had full confidence in the “wait” state at the start of each trial. The **B** matrix for sequence type was an identity matrix meaning that the agent believed a priori that the sequence of states would be stable throughout the trial. We set up the **B** matrix for the trial phase hidden state factor such that each state successively transitioned to the next state (i.e. state 1 transitioned to state 2 and so on). For the controllable **B** matrices there was one matrix for each possible report state (wait, unseen, and seen). From time step 1 - 4 the agent could only select the “wait” matrix but at time step 5 the agent could choose (i.e. via policy selection) between the “seen” and unseen “matrices”. The **C** matrix encoded the agents preference for each outcome and had a column for each time point and a row for each action (report state). There was one **C** matrix for each outcome modality. Here we only display the **C** matrix for the “correct/incorrect feedback” outcome modality associated with the report state, as it is the only outcome for which the agent had non-zero preferences. That is, the agent preferred to be “correct” at the end of each trial rather than “incorrect”. Finally, the **A** matrices were set up such that the sequence type hidden state factor had two corresponding outcome modalities, which mapped the sequence type hidden states to the internal stimulus and peripheral stimulus hidden states at level 1. To provide the model with a plausible “ignition threshold,” we lowered the precision of the **A** matrix for the “sequence type” factor by passing it through a softmax function (precision = 0.8) although we do not picture this graphically. The report hidden state factor did not map to hidden states at the level below, instead it mapped to observations that informed the agent about whether they were “correct” or “incorrect” (recall that, because of the **C** matrix, the agent wanted to receive “correct” observations and was averse to “incorrect” observations). Finally, the report hidden state factor mapped to first level language processing hidden states so that after time step 5, once the model was in a “seen” or “unseen” state, the appropriate sequence of spoken word states would be initiated at the level below (i.e. “I” “see” “a” “square”). Again, for visual simplicity the matrices displayed above have not been factorised and appear differently to how they are implemented in code.

Finally, we constructed the **C** matrix such that when the agent received feedback at time point 8 they most preferred to be correct and least preferred to be incorrect when reporting whether or not they had seen the stimulus (preference values that produced sufficient motivation for accurate reporting are depicted in figure A2). To model forced-choice behaviour, we ran a separate simulation but reduced the preference for being correct versus incorrect, making the agent less conservative and more likely to guess under conditions of weaker perceptual signals. Because Active Inference models are deterministic, we set policy precision (the confidence in policy selection denoted by β), and motor stochasticity (randomness of action selection denoted by α) to β= 2 and α= 6, thereby allowing for a plausible level of behavioural variability reflecting the agent’s relative confidence in some states over others.

## Footnotes

1 It should be noted that in the standard active inference simulation routine (spm_MDP_VB_X.m), there is an optional parameter (mdp.erp) that encodes the assumed degree of decay or attenuation in posterior beliefs across timesteps (i.e., before each period of gradient descent following each new observation). In empirical work, this parameter needs to be fit to neuronal or behavioural responses. Here we do not include this parameter (i.e., we set it to a value of 1), which entails that posterior beliefs at one timepoint fully carry over as priors for the subsequent timepoint in a trial. This is analogous to predictive coding formulations and is most appropriate for our simulations due to the fact that multiple timepoints in our simulated trials are treated as single observations (i.e., there should be no attenuation or ‘resetting’ of priors within a single stimulus presentation). It is important to note this here, however, as predicted ERPs in simulations can be affected by different values of this parameter.

## Notes

### Competing Interest Statement

The authors have declared no competing interest.

